# Neural correlates of visual object recognition in rats

**DOI:** 10.1101/2023.09.17.555183

**Authors:** Juliana Y. Rhee, César Echavarría, Ed Soucy, Joel Greenwood, Javier A. Masís, David D. Cox

## Abstract

Invariant object recognition—the ability to recognize objects across size, rotation, or context—is fundamental for making sense of a dynamic visual world. Although initially believed to be unique to primates due to its complexity, emerging evidence suggests rodents, too, can recognize objects across a range of identity-preserving transformations. Here, we describe a comprehensive pipeline for investigating visual behavior in rats, from high-throughput training to cellular resolution imaging in awake, head-fixed animals. Using this suite of tools, we demonstrate that rats excel in visual object recognition and explore potential neural pathways which may support this capacity. We leverage our optical approach to systematically profile multiple visual areas with responses to a range of stimulus types recorded in the same neurons. Primary and higher-order areas of rat visual cortex exhibit a hierarchical organization consistent with a role in visual object recognition. However, marked deviations from the functional organization of primate visual cortex suggest species-specific differences in the neural circuits underlying visual object recognition. This work reinforces the notion that rats possess sophisticated visual abilities and offers the technical foundation for their use as a powerful model to link neuronal responses to perception and behavior.

## Introduction

Generalizing across contexts is crucial for operating in a dynamic world, and for many species, vision is a critical modality for identifying perceptual regularities in the environment. There is tremendous variation in the vast number of unique retinal images a given object can cast–yet, we rapidly recognize thousands of distinct object classes with an ease that belies the computational complexity of this feat^1,2^. In primates, a series of hierarchically organized cortical areas, called the ventral visual pathway, is thought to achieve this by integrating feature-selective receptive fields of lower-level areas to form neural representations in higher-level areas that are increasingly selective for complex shapes and tolerant to identity-preserving changes in scale, position, or view^1–5^. Despite significant progress in engineering machine vision systems that solve recognition tasks at human-comparable levels^6–8^, mechanistic insight into how the brain achieves robust and ‘invariant’ object recognition remains elusive due to the limited number of animal models where this capacity can be dissected with genetic and spatial precision. Rats have been shown to perform tasks that require them to visually recognize objects across a range of contexts^9–17^, suggesting they may serve as a paradigm of complex visual behavior in which neural circuits are both genetically and optically accessible. However, the lack of methods for targeted access to neural populations in rats has limited the ability to link visual circuits to perceptual behavior.

Investigations of visual circuitry in mice^18–30^, rats^31–35^, and squirrels^36,37^ suggest that rodents have far more sophisticated visual systems than previously appreciated. The development of genetic tools and optical methods in rodents^38–41^, primarily in mice, has propelled our understanding of rodent visual physiology. As in primates, rodent visual cortex exhibits feature-selective receptive fields^25,42^ and a broadly hierarchical organization^22,23,26–28,43^. However, disparities exist between primates and rodents, such as the absence of orientation preference maps^44,45^, non-canonical collicular pathways to extrastriate areas^46^, and a shallower hierarchy^23,47^ in rodent visual cortex, possibly reflecting distinct ecological niches and inherent differences in the statistics and requirements of visual perception between species. Elucidating common and divergent principles of visual processing across species requires a comparison of both the visual circuits and the behaviors they may subserve.

Studies spanning decades^9–14,16,48–51^ have highlighted the visual behaviors and shape recognition abilities of rats. Compared to mice, rats have long been preferred in experimental settings that demand the ability to learn complex behaviors, spanning vision^9–11,13,52^, spatial navigation^53–56^, decision-making^57–60^, working memory^61–63^, and motor skill learning^64–66^, spurring developments of advanced systems to train large animal cohorts in parallel^58,63–67^. Due to the inherent variability of animal behavior, well-controlled and scalable assays are key to understanding mechanisms of perception and cognition that are shared across species. Modular, high-throughput behavior systems thus facilitate systematic investigations of core computations underlying complex behaviors such as visual object recognition. However, such systems have yet to be scaled up for visual tasks in rats.

At the neural level, tracing^34,35,68–70^ and electrophysiology^71,72^ results from early studies revealed rat visual cortex to be composed of primary sensory and putative higher-order areas. More recent multi-and single-unit recordings have described select areas of rat temporolateral cortex that may be hierarchically organized, with lateral extrastriate areas exhibiting larger receptive fields and greater tolerance to identity-preserving transformations^31–33,73^. Still, studies of rat visual circuits lag behind those in mice, particularly in the exploration of optical methods, despite increasingly available genetic tools that make these methods easier to apply^74–79^. Except for V1^45,76,80–84^, optical imaging in awake rats has proven challenging. Their larger size and strength places technical limitations on leveraging the advantages of visual experiments in rats that are awake under head-fixation. Moreover, it remains hard to access the temporolateral-most regions of rodent cortex with standard imaging approaches, and thus, higher-order visual areas proposed to play a role in visual object recognition, such as areas LI and LL, as well as parahippocampal areas implicated in visual and spatial memory^85^, remain understudied.

Here, we describe advancements in behavior and optical imaging techniques for rats that we have developed to study visual perception and neural circuits in awake animals. We devised a modular platform to automatically train large cohorts of rats on visual tasks, demonstrating their robust object recognition capacities across changes in object identity and view. We then designed a tilting two-photon microscope for imaging in awake, head-fixed rats to access the neural representations underlying these behavioral capacities. Characterizing multiple areas of temporolateral visual cortex with a range of stimulus types in the same neurons revealed a primate-like hierarchical organization in the rat consistent with a role in visual object recognition, but also non-hierarchical features better resembling mouse visual cortex. By developing methods for chronic, cellular resolution imaging in awake, head-fixed rats, our work offers insights into primary and higher-order rat visual cortex that pave the way for mechanistic studies of perception and behavior in the rat.

## Results

### High-throughput training shows robust visual object recognition

Distinct visual objects and different views of the same object can both appear as dramatic morphological differences at the retinal level. We sought to understand how animals distinguish between feature differences that indicate common object identity from those that indicate the presence of a different object altogether. One caveat to studying complex behaviors like invariant visual object recognition is that they can take many weeks of training^58,67^, which, when coupled to neural imaging, may lead to prohibitively high attrition rates. A high-throughput approach to train large cohorts (>30 animals) can overcome this challenge, as demonstrated with decision-making^58,59,86^, working memory^67^, and motor tasks^64–66^, but standard rat vision studies often employ only ∼4-6 animals per experiment^12,14,16,87^. Thus, our goal was to take existing visual paradigms^11–14,16^, and adapt them for a high-throughput behavior platform.

We developed a modular system, OpenRatBox (**Figure 1A-D, S1A**), that allowed us to run many visual behavior experiments automatically and in parallel. Using this system, we trained rats on a standard two-choice paradigm^11^, where they were required to discriminate between two target objects (objects A and B), and subsequently tested on novel images that represented either different views of the target objects or morphed versions of the objects presented at familiar views (**Figure 1A**). Water-restricted rats were trained to initiate a trial by licking a central sensor, wait for a stimulus to appear on a screen, then report the object’s identity by licking a left or right response port (**Figure 1B**). Correct choices were rewarded, and incorrect choices were punished (see Methods). Rats successfully performed several hundred trials per day (**Figure S1B**), and results were highly reproducible across many animals and boxes (**Figure 1E-F**, n=48/56 reached criterion performance of >70% correct across 12.6 ± 7.6 sessions). Once rats reached criterion performance of 70% accuracy (see Methods), we tested them with new images that corresponded to either identity-changing (**Figure 1G-H**) or identity-preserving transformations (**Figure 1I-K**).

**Figure 1.**
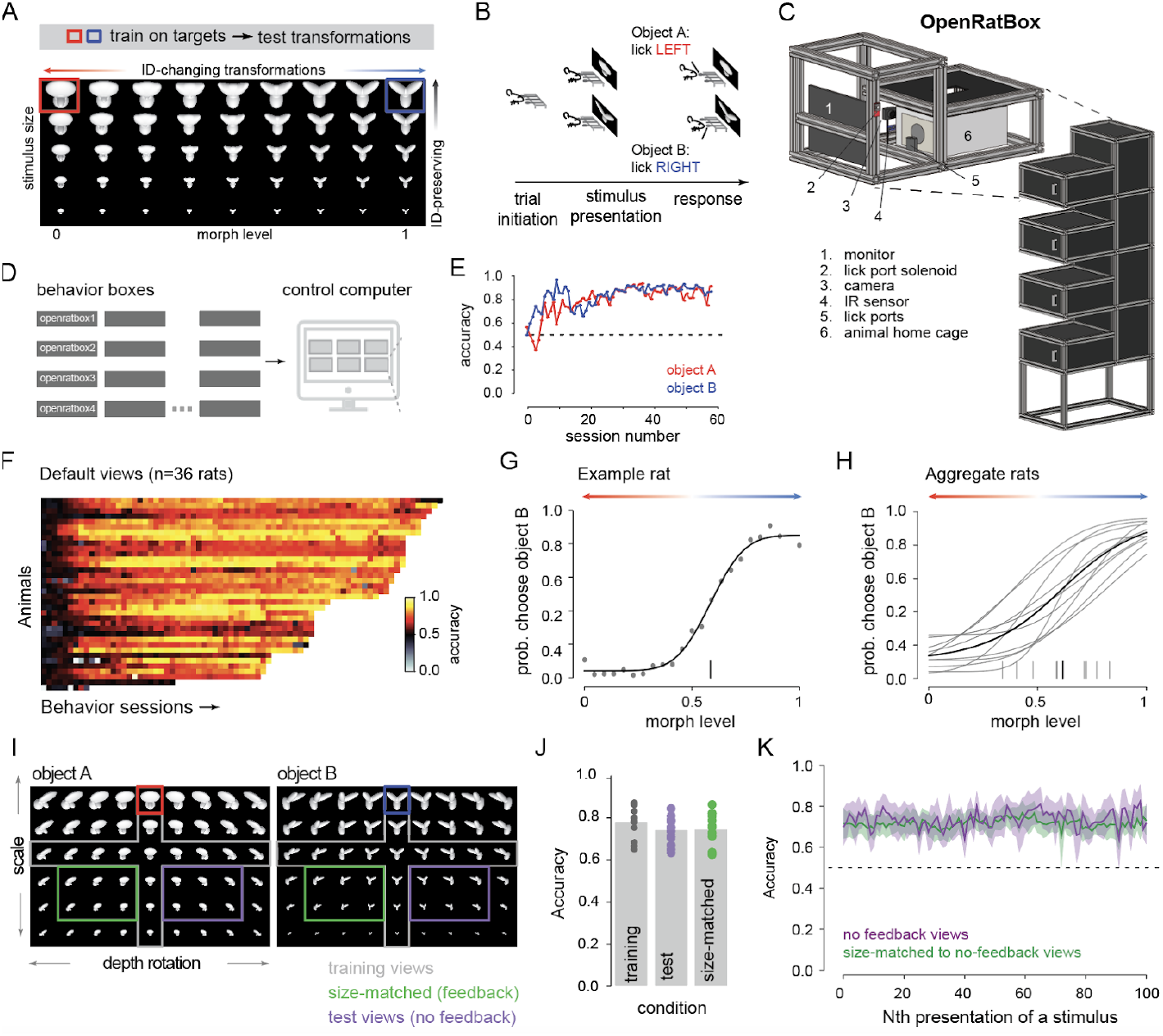
Automated, high-throughput training of visual behavior in rats. **A**, Rats were first trained to recognize two target objects, A and B (red and blue boxes). Rats that passed criterion (see Methods) were then tested on novel images that changed object identity (‘ID-changing transformations’) or object view (‘ID-preserving’; only size shown, but rotation also tested as in (I)). **B**, Schematic of trial structure (adapted from Zoccolan et al.^11^). **C**, Schematic of OpenRatBox. Four training boxes are stacked on the right, with a zoomed-in view of one box on the left. **D**, Control logic of OpenRatBox (experiments run with MWorks). Each box runs a server for a specified experiment, and one primary computer connects via a client that controls independent boxes in parallel. **E**, Training time course for an example rat. Red, accuracy for object A. Blue, accuracy for object B. **F**, Average accuracy by session for a cohort of rats (n=36 rats). Each square represents accuracy in one session per animal. **G**, Psychometric curve for an example rat on identity-changing transformations. Circles, data. Line, Gaussian fit (see Methods). Vertical bar, fit point of subjective equality (PSE). **H**, Psychometric curves for a subset of trained rats tested on identity-changing transformations (n=10 rats). Gray, individual rats. Black, average across rats. Vertical bars, points of subjective equality (PSE). **I**, Identity-preserving transformations for each target object (red and blue boxes). Gray, training views (see Methods). Purple, example test views for which feedback was never provided. Green, size-matched views of the test views, but for which feedback was provided. **J**, Average accuracy on identity-preserving transformations for training (gray), no-feedback (purple), and acuity-matched with feedback (green) views. Each dot represents one animal (n=13 rats). **K**, Average accuracy on identity-preserving transformations as a function of the N-th presentation of a given view for no-feedback (purple) and acuity-matched feedback conditions (green). Shading, standard deviation, s.d.

The identity-changing transformations assessed how well rats discriminated between the two target objects. Morphs, spanning morphological variations in identity from level 0 (0% B and 100% A) to 1 (100% B and 0% A), were presented on a small fraction of probe trials (<15%) on which no feedback was provided, and the category boundary between the targets was determined with a psychometric curve^88^ relating the animal’s choices to morph levels (see Methods). Rats’ perceptual choices were well-aligned with the stimulus space (**Figure 1G**). As morphs became more ‘B’-like, rats increasingly reported object B, and vice versa, with the point of subjective equality (PSE, where reports of A and B were equally likely) slightly closer to B than A (**Figure 1H**, 0.65±0.16, mean±s.d. across n=10 rats).

How precisely rats differentiated between the objects was captured by the difference threshold, or the minimum morph difference perceived 50% of the time (just-noticeable difference, JND, half the morph level spanning 25% and 75% ‘B’ choices), where larger values reflect flatter, less discriminative curves and smaller values correspond to steeper curves and sharper category boundaries. Rats perceived differences of ∼20% between morphs (0.21±0.09 morph levels, mean±s.d., n=10 rats), underscoring their robust discriminatory capacity, which we later leveraged to compare to the discriminatory capacity present in the activity of neural populations (see below).

We then assessed how well rats recognized the target objects across identity-preserving transformations. Each object was presented at different scales and rotations (**Figure 1I**), yielding greater pixel-level, and corresponding retinal level, differences between two views of the same object than two objects shown in the same view (see Methods). Since rats could, in theory, use feedback to memorize the correct response for each image, we withheld feedback from a subset of views to test spontaneous generalization (purple, **Figure 1I**). However, low accuracy on these no-feedback views could reflect poor generalization or acuity limits, especially for small stimulus sizes, so we also compared performance on corresponding size-matched views for which feedback was provided (green, **Figure 1I**). All trained rats generalized to novel views, maintaining high accuracy across all conditions (**Figure 1J**; default: 0.78±0.07, no-feedback: 0.74±0.06, size-matched: 0.74±0.08, mean± s.d., n=13 rats), consistent with previous studies^11^. Importantly, accuracy was not influenced by the number of exposures to a given view (**Figure 1K**), demonstrating that generalization did not depend on trial history, but could be an intrinsic feature of the underlying neural representations rather than emerging only with learning.

Although both identity-preserving and identity-varying transformations involve large changes in the images falling on the retina, our behavioral results demonstrate that rats accurately distinguish between changes that alter object identity and those that preserve it. Importantly, despite that moving visual objects provide higher salience to rodents^32,89^, our trained rats exhibited robust generalization and discrimination performance even with static images, similar to paradigms used in primate studies of visual object recognition. Having empirically determined both the discriminability and generalizability of our object stimuli at the behavioral level, we next sought to probe the neural representations that might support these behaviors in rats.

### A complete pipeline for neural imaging in awake, head-fixed rats

Imaging in rats has been challenging due to their larger brain size—requiring a bigger field-of-view (FOV)—and significant strength from their larger body size^90^, which can lead to motion-induced image distortions or even implant detachment from the skull. To further complicate matters, higher-order visual areas in the rat proposed to be analogous to the primate ventral visual stream^31,32,91,92^, including areas LI and LL, are positioned on the side of the animal’s head. These areas are difficult to access using standard, upright microscopes or head-mounted mini-scopes^83,90,93,94^, and thus, little is known about their functional properties. To overcome these hurdles, we developed a pipeline for chronic access to lateral visual cortex in awake, head-fixed rats (**Figure 2A, S2A**).

**Figure 2.**
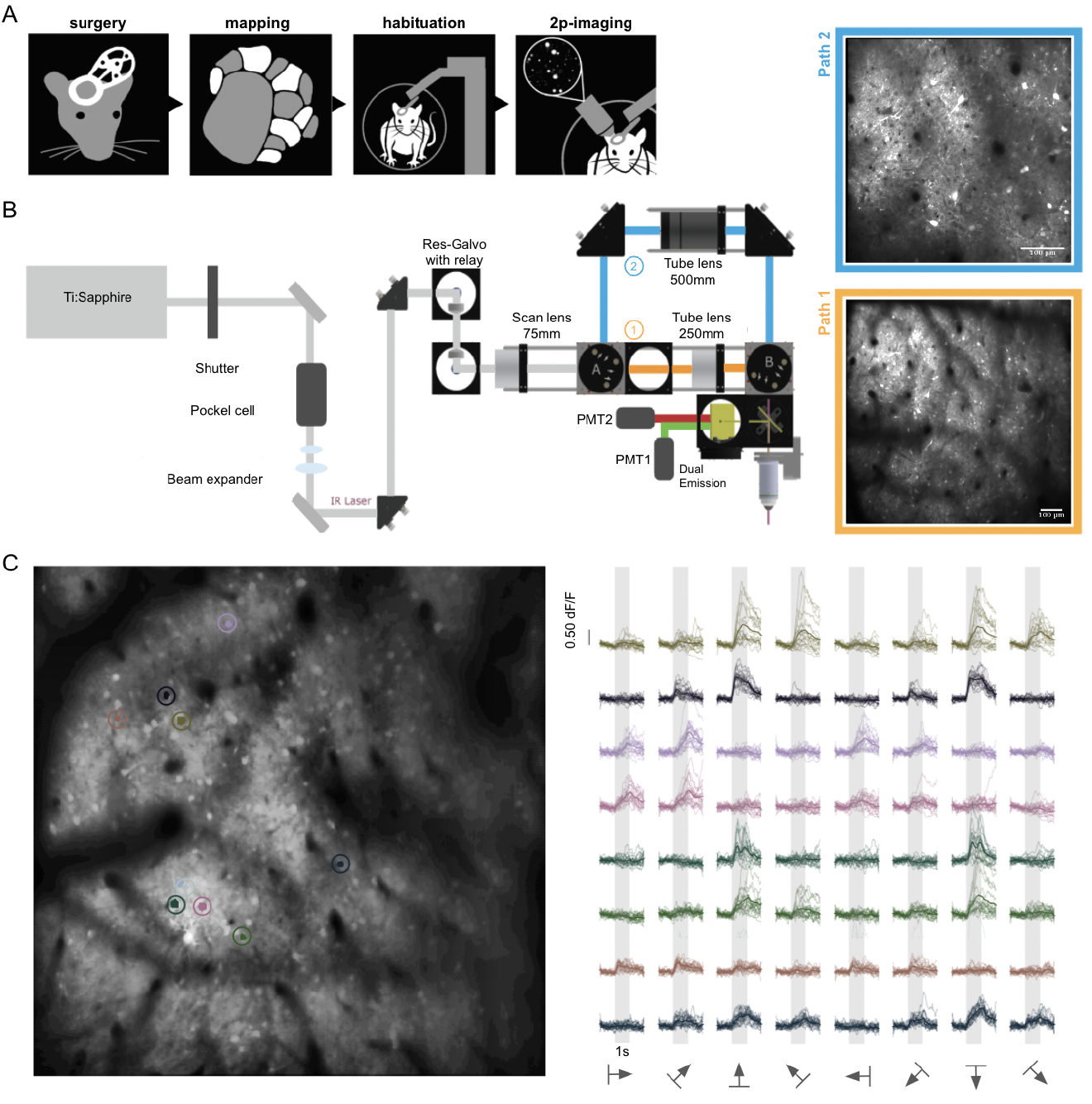
A standardized pipeline for optical imaging in awake, head-fixed rats. **A**, Schematic of the workflow for each rat in our imaging pipeline. **B**, Schematic of the tiltable imaging system with dual-channel two-photon microscopy. Two-photon beam paths provide either a standard-scale (cyan) or a large-scale (orange) FOV half the zoom of the standard mode. Both modes provide cellular resolution at the same acquisition rate. *Top*: Example standard FOV (500 μm x 500 μm) showing neurons labeled with GCaMP7f. *Bottom*: Example large FOV (1 mm x 1 mm). Scale bars, 100 μm. **C**, *Left*: Example two-photon large FOV of GCaMP7f fluorescence in V1. *Right*: Example time courses in response to drifting gratings for the 8 neurons circled with corresponding colors on the left. Traces are stimulus-aligned responses for 8 directions of gratings presented at the spatial frequency, speed, and size combination that elicited the cell’s maximum response. Thin lines, individual trials. Thick lines, mean response. Gray bars, stimulus period (1 sec).

To keep the rat’s body in a natural, resting position and simplify the presentation of visual stimuli, we rotated the microscope objective relative to the animal via a tilting, dual-channel two-photon microscope that allowed access to virtually any imaging plane around the animal’s head, while maintaining over 180º of unobstructed viewing angle (**Figure 2B, Figure S2E-F**). Two optical paths provided FOV sizes of ∼500x500-1000 µm^2^ to 1x1-2 mm^2^ (**Figure 2B**, top and bottom insets), each capable of cellular resolution imaging of neural responses (**Figure 2C**). Dual-channel epifluorescence paths facilitated vasculature and retinotopic mapping of targeted sites (**Figure S2E**,**G**).

A key challenge was optimizing surgical and habituation procedures for extended head-fixation with angled head plates in awake rats and clear windows lasting several weeks (see Methods, **Figure S2A**). Our methods enabled robust implants and stable imaging at >4 weeks post-surgery (**Figure S2A**). Custom titanium head plates were implanted to align the cranial window to the imaging plane (30 degrees, see Methods), and mated with a custom kinematic mount (**Figure S2C-D**) that provided sufficient spatial precision for returning to the same cells across days (**Figure S2H**). These methods thus enabled us to characterize populations across multiple visual areas to shed light on the network of neural circuitry that supports complex visual behavior.

### Optical mapping of rat visual cortex

To optically map rat visual areas, we relied on multi-site injections of a genetically-encoded calcium indicator^95^ and a large (5 mm diameter) window placed over the posterior-temporal edge of rat cortex, targeting areas V1, LM, and LI (see Methods). We developed a tilting tandem-lens epifluorescence macroscope^96^ and performed widefield mapping of the entire window (**Figure 3A**) with a standard, cycling bar paradigm presented on a large monitor centered on the eye contralateral to the window^18,20,97–99^ (see Methods). Rat visual cortex exhibited smooth retinotopic maps delineating primary and higher-order visual areas (**Figure 3B-C**), as observed in other species mapped with optical imaging^18,20,98,100,101^, and optical maps were qualitatively similar to electrophysiological estimates of areas in rat visual cortex^71,72,102^. Fast optical mapping of the entire cranial window thus allowed us to reliably target V1, LM, and LI for two-photon imaging.

**Figure 3.**
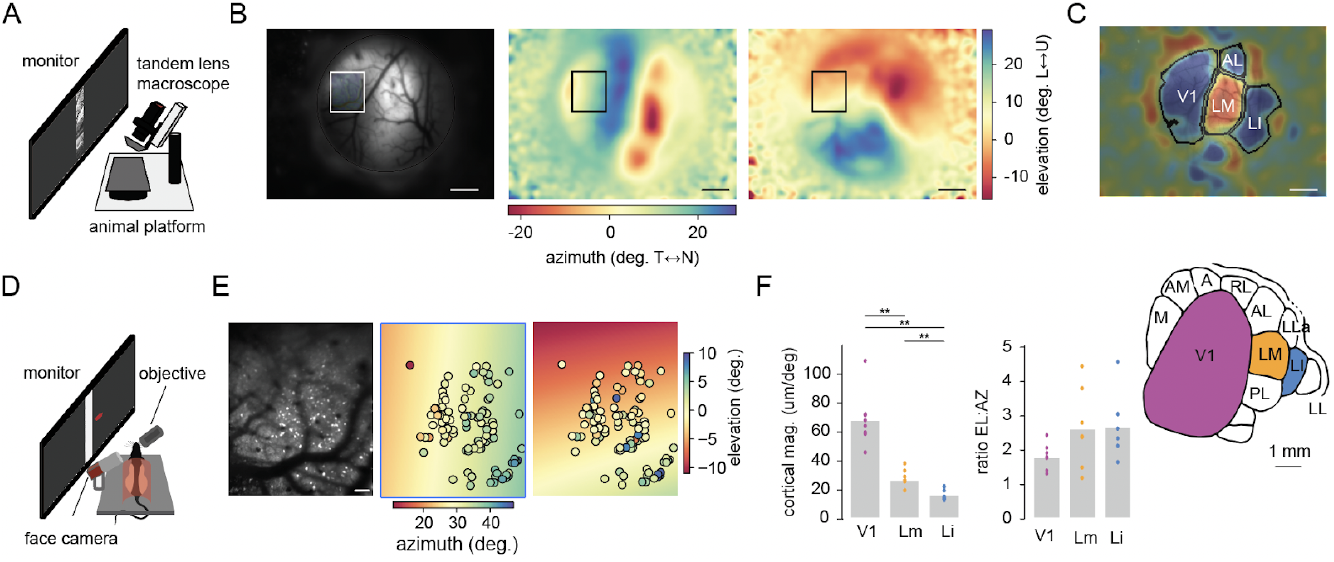
Retinotopic organization of rat visual cortex. **A**, Schematic of the tilting tandem-lens macroscope setup for widefield mapping. **B**, Widefield maps. *Left*: Epifluorescence image of the cranial window. Outlined box corresponds to a V1 two-photon imaging site. Pseudo-colored images from phase-encoded mapping of retinotopic preference along azimuth (middle) and elevation (right) (see Methods). T=temporal side of visual field, N=nasal side, L=lower visual field, U=upper visual field. Scale bar, 1 mm. **C**, Visual field sign maps calculated from the azimuth and elevation maps shown in **B. D**, Schematic of the two-photon imaging platform. An IR camera captured high-resolution video of the animal’s face synced with neural data acquisition. **E**, Two-photon maps. Max-projection of GCaMP fluorescence for the site outlined in **B** (left) and pseudo-colored retinotopic maps for azimuth (middle) and elevation (right). Circles, cell body locations colored by retinotopic preference (Methods). Scale bar, 100 μm. **F**, *Left*: Average cortical magnification for FOVs in V1, LM and LI. *Right*: Ratio of cortical magnification along elevation (EL) versus azimuth (AZ). Inset, visual areas targeted in this study (adapted from Olavarria et al.^34^ and Sereno et al.^102^). Bars, mean across imaging sites. Circles, average for each FOV. Colors correspond to areas shown in the brain inset. See Methods for statistical comparisons.

With the same stimulus paradigm as used for widefield mapping, we measured retinotopy in two-photon FOVs (**Figure 3D-E**). Cortical magnification, or the extent of cortex representing a given portion of the visual field, decreased along V1, LM, and LI (**Figure 3F**), consistent with their decreasing surface areas^71,102^. In contrast to the radial anisotropy of primate visual cortex, where cortical magnification decreases with eccentricity^103,104^, rat visual cortex exhibited anisotropy in azimuth and elevation, with a two-fold greater cortical magnification along elevation (**Figure 3F**), similar to the anisotropic expansion seen in mouse visual cortex^44,98,99,105,106^.

### Single neuron response profiles distinguish primary and higher-order visual areas

In primate ventral visual cortex, decades of studies probing many stimulus types and transformations have established a rich view of area-specific characterizations, where the change in response profiles from one area to the next are thought to support a progression of increasing selectivity for complex objects and greater tolerance across views. For example, bigger receptive fields provide access to larger portions of the visual field, which may facilitate scale and position tolerance^3,107,108^, while reduced responsiveness to simple edges in favor of more complex feature conjunctions may support greater object selectivity^2,109–113^. In contrast, most studies of rodent visual cortex have relied on limited stimulus sets (though see^21,114^), uneven coverage or biased sampling of neurons, or both, making direct comparisons across areas and stimulus types challenging. A key advantage of our experimental setup is that it allowed us to systematically profile multiple visual areas with a broad range of stimulus types in the same neurons (**Figure 4**). Specifically, we estimated receptive fields by flashing small gratings across the visual field **(Figure 4A-C**), and measured axis and direction tuning with drifting square-wave gratings (**Figure 4D-F**). Finally, to see how changes in tuning across visual areas from simple features to complex shapes might support the object recognition capacities observed in our trained animals, we characterized responses to the same object stimuli tested in our behavior task (**Figure 4G-J**).

**Figure 4.**
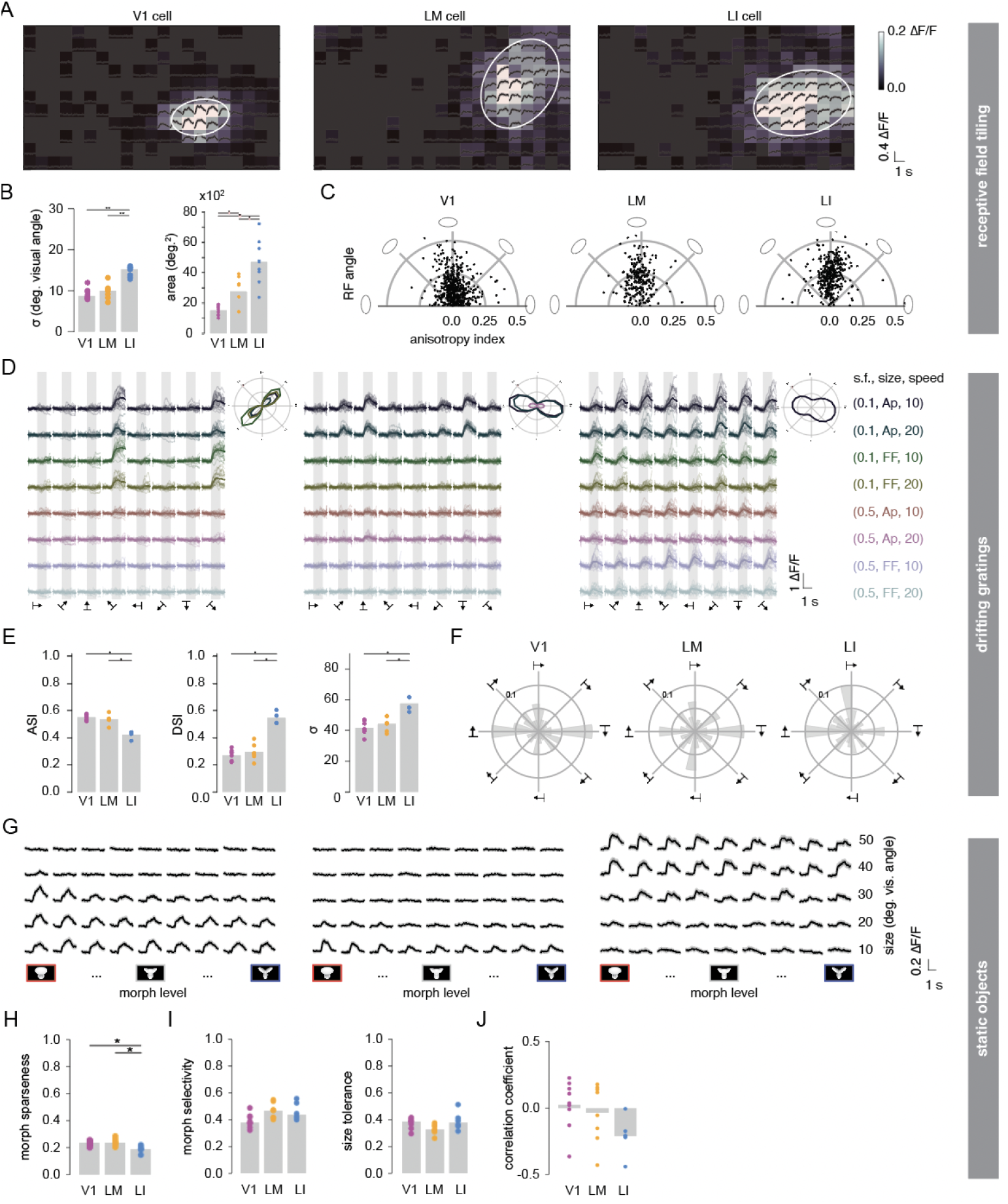
Single neuron responses across primary and higher-order visual areas. **A**, Receptive fields of example neurons from V1 (left), LM (middle), and LI (right). Traces, mean time courses 1 sec from stimulus onset for the small gratings tilling the visual field, flashed for 0.5 s at each stimulated location. Heat map, mean response. Ellipse, fit receptive field at full-width-half-maximum (FWHM). **B**, *Left*: Average receptive field sizes. *Right*: Population receptive field sizes. Each dot represents one FOV. Bars, mean across sites. **C**, Distribution of receptive field angle and anisotropy by visual area. Each dot represents one cell. Theta axis, receptive field angle (0° is parallel to the horizontal plane). Radial axis, anisotropy index, defined as the difference between the major and minor axes of the fit ellipse, divided by their sum (0 is perfectly isotropic, or a circle, 1 is perfectly anisotropic, or a line). **D**, Responses to drifting gratings from an example cell in each visual area. Thin lines, individual trials. Thick lines, trial means. Colors, distinct combinations of spatial frequency, size, and speed (Ap=apertured, FF=full-field). Columns represent drifting direction. Polar plot insets at the upper right for each cell show corresponding direction tuning curves that pass goodness-of-fit criteria (see Methods). **E**, Axis-selectivity index (ASI, left), direction-selectivity index (DSI, middle), and tuning width (sigma σ, right) estimated from fit direction tuning curves (see Methods). Each dot represents the median across cells for one imaging site. Bars, mean across sites. **F**, Polar plots showing normalized distributions (fraction of fit cells) of preferred direction of motion for each visual area. **G**, Responses to objects for example neurons from V1 (left), LM (middle), and LI (right). 5 stimulus sizes (identity-preserving) and 9 morph levels (identity-changing) were tested, in addition to 5 full-field, grayscale stimuli matched in overall luminance to each stimulus size (see Methods). Traces, mean±s.e. time courses 1 sec before and after the 1 sec stimulus presentation. **H**, Lifetime sparseness of non-luminance-preferring cells (see Methods). Dots, median across cells for one imaging site. **I**, Morph selectivity (left) and size tolerance (right) calculated at each cell’s best size or best morph, respectively. **J**, Pearson’s correlation coefficient between morph selectivity and size tolerance for each site.

Rat ventral visual areas exhibited a broad, primate-like hierarchical organization. From primary to higher-order visual areas, receptive field sizes increased (**Figure 4B**), as previously observed in rodent extrastriate areas^22,23,31,32,42,115^, while orientation selectivity decreased, direction tuning curves broadened^18,32^ (**Figure 4D-E**), and fewer cells had well-fit direction tuning curves (though direction selectivity was higher among LI cells that were direction-tuned; fraction of well-fit cells, V1: 0.50±0.10, n=8 FOVs, LM: 0.50±0.13, n=7 FOVs, LI: 0.27±0.06, n=4 FOVs). These results are consistent with evidence from primates^2,3,111,116^ that visual areas form a hierarchical network characterized by features such as increasing receptive field sizes and decreasing preference for simple features.

However, we also observed marked deviations from the primate visual system in all three areas that resembled observations of mouse V1, including fine-scale retinotopic scatter at the level of single neurons^44,105^ despite smooth widefield maps (**Figure S3A-C**, and see **Figure 3E**), salt-and-pepper feature maps without strong spatial correlations^44,45,117^ (**Figure S3D**), and consistent with widefield measurements, anisotropic receptive fields with cortical magnification expanded in elevation^98,105,106^ (**Figure 4C**, mean±s.d. ratio of major to minor axis, V1: 1.43±0.21, n=9 sites, LM: 1.49±0.08, n=7 sites, LI: 1.59±0.19, n=9 sites). The observed anisotropy could not be explained by spherical distortions from a flat monitor: applying the inverse transformation of the spherical correction to measured receptive field maps (see Methods) resulted in quantitatively similar receptive field metrics (**Figure S4**). Rather, each cell’s receptive field appeared to correspond to a smaller portion of the visual field along elevation relative to azimuth, suggesting that rodents may have enhanced resolution along the vertical axis of view.

Despite that some properties of rat extrastriate areas resembled features of mouse visual cortex, our behaving rats nevertheless showed robust invariant object recognition performance at levels higher than mice trained on moving stimuli^28^, suggesting the existence of neural representations supporting these behaviors.

We thus probed neural responses to a subset of the same stimuli used in our behavioral paradigm, to link single neuron metrics of selectivity and tolerance to stimuli with known perceptual responses (**Figure 4G-J**).

In nonhuman primates, both selectivity for complex shapes and tolerance to identity-preserving transformations increase in a balanced manner such that lifetime sparseness, or the specificity of neural responses to object stimuli, is preserved across the ventral stream^4,118^. This is thought to occur as a trade-off at the level of single-units exhibiting a negative correlation between selectivity and tolerance, as seen in the latest stage of the primate ventral pathway, inferotemporal cortex, where units more selective to specific objects are less tolerant to transformations, while highly tolerant units are less selective^119^. We therefore estimated similar metrics across rat visual cortex. Lifetime sparseness^120,121^ (see Methods), was comparable across visual areas for the object stimuli tested, though slightly lower in LI (**Figure 4H**; V1: 0.22±0.02 across n=9 FOVs, LM: 0.23±0.0 across n=8 FOVs, LI: 0.18±0.02 across n=6 FOVs, mean±s.d.). This difference could be due to lower selectivity for morphs, greater tolerance to changes in scale, or a combination of both.

To further examine the interplay between selectivity and tolerance in rat extrastriate areas, we calculated a morph selectivity index and size tolerance index for each neuron, and estimated the trade-off between the two^119^. Morph selectivity captures how much a neuron responds to the various morphs presented at the neuron’s preferred stimulus size, while size tolerance measures how much a neuron’s response to its preferred morph changes across scale. In contrast to the primate ventral pathway, we did not observe significant differences between areas in single neuron metrics of morph selectivity or size tolerance (**Figure 4I**), and although LI sites tended to exhibit more negative correlation coefficients between selectivity and tolerance (**Figure 4J**), areal differences were not statistically significant.

Altogether, though single neuron response profiles across lateral visual cortex were broadly consistent with a primate-like hierarchical pathway, many features, not only of rat V1, but also LM and LI, were not, with some resembling mouse V1, suggesting both shared and species-specific features of visual processing in rodents and primates.

### Population representations support increased generalization in higher-order visual cortex

Though single-neuron metrics of object selectivity and tolerance revealed far more subtle area differences than the dramatic specializations seen across the primate ventral stream, certain feature-selectivities in rat higher-order areas, such as reduced preference for simple gratings stimuli and increasing receptive field sizes, suggested that population-level representations might nonetheless support the recognition capacities observed in our behavior paradigm. Importantly, rats performing the visual object recognition task accurately classified novel transformations in the absence of feedback (**Figure 1**), suggesting that neural representations supporting generalization are inherent to the circuitry of their visual systems.

One biologically plausible scheme for how neural populations perform computations enabling visual object recognition is a thresholded sum taken over weighted synapses to transform representations^4,122^. We thus explored neural population activity with linear classification models (linear support vector machines) and estimated discriminability and generalization in primary and higher-order visual areas. We first trained linear classifiers to discriminate between target objects A and B using population responses to the same stimuli as used for behavior. We then tested these classifiers on identity-changing morphs or identity-preserving changes in stimulus scale, matching the tests performed by our trained animals (**Figure 1A**). If discriminability between objects is high, the classifier should report object A when A is shown, and vice versa for object B (**Figure 5A**), and likewise, if neural responses to morphs reflect both the stimulus space and animals’ perception, classifiers should categorize neural representations as object B more often for morphs closer to B than to A, and vice versa.

**Figure 5.**
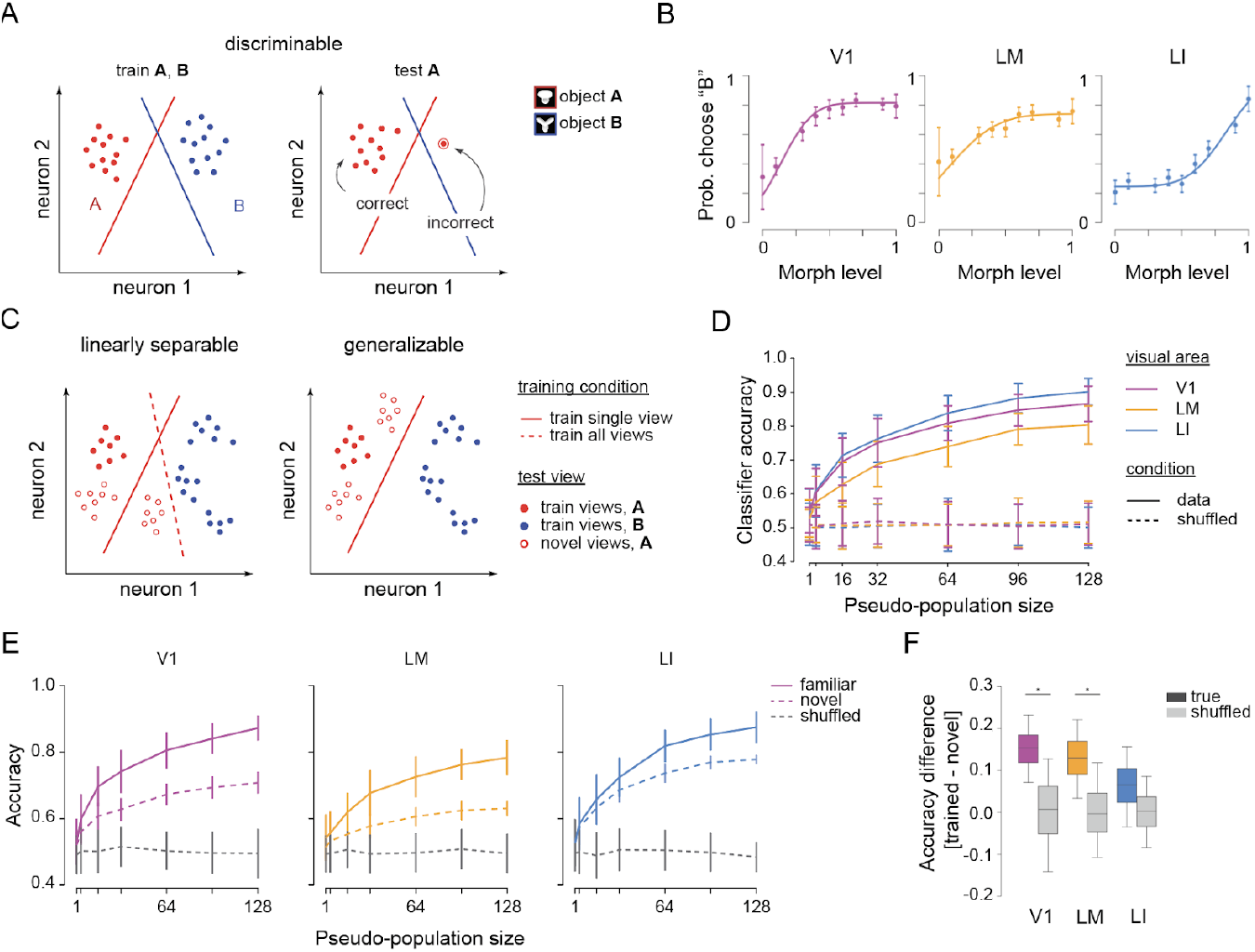
Linear separability and generalization of neural representations. **A**, Schematic (adapted from Rust et al.^4^) illustrating discriminability measured with linear classifiers (see Methods). For each object, the optimal linear hyperplane (line) separates responses to one object from responses to all others. Each dot represents the neural population vector for one trial. A subset of trials are used to find each hyperplane (left), then discriminability is measured using the remaining trials (right). The percent of response vectors falling on the correct side of the hyperplane (left side, if A is shown) as opposed to the incorrect side (encircled dot) estimates the similarity between a given response and other responses to a given object. **B**, Population neurometric curves. Linear classifiers were first trained on the target objects, then tested on the intermediate morphs. Probabilities of choosing ‘B’ at morph levels 0 and 1 reflect discrimination accuracy for the target objects. Circles and bars, mean±s.d., n=500 bootstrap iterations. **C**, Schematic illustrating linear separability and generalization (adapted from Rust et al.^4^). Training on one stimulus condition (left) may fail to separate response vectors correctly (solid red line), *i*.*e*., fail to generalize to novel conditions, while training on all transformations simultaneously may identify a better boundary (red dashed line). In contrast, training on a single view might be sufficient for populations to generalize to novel conditions (right). **D**, Classifier accuracy when trained and tested on all stimulus sizes as a function of population size. **E**, Classifier accuracy for trained (solid) and novel (dotted) test conditions. Gray, accuracy when object labels were shuffled. **F**, Generalization capacity, defined as the difference between accuracy on trained versus novel conditions. Gray, generalization capacity when ‘trained’ and ‘novel’ labels were shuffled.

Classifier performance indicated good discrimination, with high decoding accuracy for each object (**Figure 5B**). Similar to the psychometric curves fit from behavioral responses (**Figure 1G-H**), we fit neurometric curves for each visual area from classifier responses to test trials (**Figure 5B**), and estimated a ‘neural PSE’ (point of subjective equality) as the morph level at which objects A and B were equally likely to be selected. V1 and LM classifiers were biased toward object A, while LI classifiers exhibited a bias toward object B (V1: 0.14±0.04, LM: 0.08±0.04, LI: 0.83±0.05, mean±s.d., i=500 iterations), the same direction as the slight behavioral biases we observed (**Figure 1H**). The biases in neurometric curves were not simply due to a lack of cells preferring one of the objects in a given visual area, since all three areas contained both cells that strongly preferred object A and cells that strongly preferred object B (**Figure S5**, see Methods), and may instead reflect nonuniform feature selectivity across visual areas.

LI’s larger receptive fields might impair discriminability such that LI classifiers require a greater difference in morph level to achieve a given level of discriminability, compared to V1 or LM classifiers. To explore this idea, we estimated a neural difference threshold, analogous to our behaviorally-derived difference thresholds, which estimated that rats perceive morph differences of about 20% (**Figure 1G-H**). The neural threshold was about half the behaviorally-perceived threshold in all visual areas (just-noticeable difference, JND, in V1: 12±7%, LM: 13±7%, LI: 13±8%, mean±s.d., i=500 iterations), suggesting these modestly-sized neural populations can, at least in theory, discriminate finer differences than indicated by behavior output.

Our trained rats also showed robust generalization across morphological changes that preserved rather than altered each object’s identity (**Figure 1I-K**). To test how this may be implemented in rat visual cortex, we compared the ability of neural populations in each area to generalize across changes in scale by testing linear classifiers in two different ways^4^. Under the first regime, simultaneously training discrimination at all scales and testing on trials excluded from training estimates linear separability in a way that does not rely on which reference image was used for training. However, linear separability does not necessarily mean good generalization^4,31^ (left, **Figure 5C**). Thus, in the second regime, training on a single reference (*i*.*e*., at a one size) and testing novel conditions may reveal linearly separable populations that fail to generalize (left, **Figure 5C**), while successful generalization suggests representations based on scale-invariant features (right, **Figure 5C**).

When we trained classifiers to categorize the target objects using all stimulus scales, decoding accuracy was comparable between V1, LM, and LI (**Figure 5D**). In contrast to the primate ventral stream^4,118^, where the ability to discriminate between objects is thought to be supported by an increasing linear separability of the neural representations corresponding to each object across the visual hierarchy^2,4,123^, under our conditions, linear separability of object representations in the rat may be relatively comparable between primary and higher-order visual areas.

To test generalization across changes in scale, we trained classifiers with images at a single stimulus size, then tested decoding accuracy at the remaining stimulus sizes excluded from training. V1 and LM classifiers performed significantly better on trained than novel stimulus sizes, while LI classifiers performed well on both (**Figure 5E**, accuracy on trained minus novel, V1:14.6±4.8%, LM: 14.5±5.2%, Li: 8.0±6.0%, mean±s.d. across i=500 iterations for population sizes of n=128 cells), suggesting that higher-order areas may be more robust to changes in scale. We computed a generalization index as the difference between test scores on trained and novel conditions: small values indicate good generalization, while large values represent poor generalization. We observed a trend towards larger generalization values in LI relative to LM and V1, but it was not statistically significant (**Figure 5F**, permutation test shuffling area labels).

Area LI’s unique robustness to changes in scale could be due to receptive field size, where larger receptive fields confer tolerance across scale by accessing more of the visual field (**Figure 4B**). To test this idea, we subsampled cells by matching their receptive field sizes one-to-one across the three areas, such that every cell with a given receptive field size in one area had a corresponding cell matched in receptive field size in the other areas (**Figure S6C**). For LM decoders, test accuracies on novel and trained stimulus sizes became statistically indistinguishable (**Figure S6D**, bottom), suggesting a partial role for receptive fields in supporting scale tolerance. However, matching receptive field sizes had no effect for V1 or LI decoders (**Figure S6D-E**; permutation, p<0.05; mean difference in accuracy ±s.d. for matched vs. non-matched populations, V1: 14±4.4% vs. 13.6±5.3%, LM: 6.5±5.5% vs. 12.9±5.4%, LI: 6.5±5.7% vs. 6.5±6.4%, i=500 iterations, n=96 cells), suggesting that receptive field size alone does not account for generalization, consistent with previous studies that did not find tolerance capacity to depend on receptive field size^31,32,73^. Rather, the format of object representations, as indicated by their linear separability, appears to confer view tolerance to areas V1, LM, and LI.

Our results suggest that rat lateral extrastriate areas can support generalization in the absence of learning or prior familiarity, consistent with findings in anesthetized^113,124,125^ or passively viewing monkeys^4,118,126–128^ and anesthetized rats^31^. Under our stimulus conditions, behavioral stimulus tolerances are likely not coded at the level of a single cell, but rather, appear distributed across a network of cells with differing degrees of response selectivity and generalization.

## Discussion

Robust object recognition maintains object cohesion amidst changing contexts, yet understanding its neural basis remains a challenge. Rodents are valuable models due to their genetic accessibility and experimental tractability, offering an inroad to link neural response properties to behaviors. Rats, in particular, are a powerful system to explore perceptual mechanisms: like primates, rats display sophisticated visual behaviors—both trained and untrained—but like mice, they are amenable to genetic circuit dissection and optical imaging approaches. Until now, methods for cellular resolution imaging in awake, head-fixed rats have been limited, relying on trained voluntary head-fixation^81^ or single-photon imaging in freely-moving animals^76,93,129–131^. We created a high-throughput visual behavior platform and a pipeline for cellular resolution imaging in awake, head-fixed rats to investigate neural correlates of invariant object recognition in rats.

Most rodent vision studies have focused on early processing stages, from retina to primary visual cortex. Our study contributes to a growing body of work dissecting the functional logic of rodent extrastriate cortex^18,21,22,27–29,31–33,114,132–144^. In particular, neural correlates in lateral areas LI and LL have remained elusive. Given the impressive visual behaviors of rats, electrophysiology studies have targeted these areas for single-unit characterizations of object responses^31,32^, identifying a candidate network analogous to the primate ventral pathway. Our recordings confirm and expand on these studies with optical imaging in awake rats, providing an important path toward spatiotemporal investigations of genetically identifiable neural populations in rats. We find that rat primary and lateral visual areas exhibit a functional hierarchy, broadly consistent with previous single-unit observations, with increasing receptive field sizes, decreasing selectivity for simple gratings, and robust generalization in higher-order areas, suggesting shared principles of visual object recognition between rodents and primates.

Still, rat visual cortex exhibited properties dramatically distinct from primate visual cortex. Our approach allowed us to explore the fine-scale retinotopy and spatial organization of neural populations, revealing mouse-like anisotropies in visual field representation^18,20,24,44,97–99,105,145^ and salt-and-pepper feature maps^45,117,146,147^ across primary and higher-order areas. While we observed robust generalization in LI populations, differences in object response metrics were otherwise subtle, consistent with prior descriptions of this area^31,32^, suggesting a loose, rather than one-to-one, mapping to the primate ventral stream. While training could pronounce object tuning differences in rat lateral cortex, specializations across the primate ventral stream have been described in anesthetized^113,124,125^ or passively viewing^4,118,126–128^ monkeys, and direct comparisons of inferotemporal cortex responses in trained and naive monkeys did not find differences^4,118,122,148^. Nevertheless, neural responses can be shaped by familiarity^149–156^, and future studies tracking neural activity during learning with methods developed here may uncover experience-dependent tuning in rats. The differences we observe between rat and primate visual systems likely reflect species-specific specializations in pathways supporting visual object recognition^43–46,98,105,145^, underscoring how cross-species comparisons of naive responses offer insight into shared and divergent properties of potentially analogous brain regions.

### Rodent models of invariant object recognition

Robust perceptual generalization, observed in our behavior results and other studies^11,13,14,16^, suggests primate-like ventral stream computations in the rodent visual system. In response to morphological transformations corresponding to identity changes, LI classifiers resembled animals’ behavior biases toward the same object, while V1 and LM classifiers displayed a bias toward the other object. Since recordings were from naive rats, the observed biases in neural representations may reflect intrinsic differences in feature selectivities between lower-and higher-order areas.

Across identity-preserving transformations, classification performance of LI populations also mirrored that of trained rats, performing well on both trained and novel conditions, unlike V1 and LM populations which showed significant performance drops. Our results using optical methods are consistent with single- and multi-unit electrophysiological studies in rats^31,32,73^, which observed better generalization capacity in higher-order areas, particularly areas LI and LL. Interestingly, a recent study in mice using moving stimuli found that, in contrast to rats, areas LM and AL exhibit the strongest generalization^28^. These results suggest there may be subtle yet consequential differences in how the visual systems of rats and mice support object perception. Importantly, optical imaging in rats–a viable alternative behavioral model to nonhuman primates–provides access to the fine-scale physical relationships between single cells within and across areas, ultimately allowing a better understanding of how spatial organization supports neural function for visual behavior.

The platform and methodology we developed here for rats offers a powerful inroad for comparative rodent vision studies, which until now have been dominated by studies of visual physiology in mice^26,27,157^. Despite their capacity for genetic manipulation, mice possess relatively poor visual acuity^49^ and exhibit inferior performance on many behavioral tasks^28,54,158–162^ compared to rats, suggesting potential limitations in their utility as a singular model for the mechanistic study of visual perception. While performance differences between mice and rats in visual tasks could be methodological, they likely also reflect species-specific differences. Rats are crepuscular^163^, are well-established to act as predators (including of mice^164–166^), have double the visual acuity of mice^49^, and have divergent extrastriate area organization^54,99,102,161,167,168^. Other, arboreal rodents, such as squirrels, are diurnal species that also display keen visual senses, though even less is known about their visual cortex beyond V1^37,169,170^. Future studies may reveal how visual systems reflect distinct sensory requirements and adaptations to specific ecological niches, which is important to consider in the context of both laboratory and naturalistic behaviors.

Historically, the study of perception has been restricted to nonhuman primates, due to the similarity of their visual systems to our own, as well as their impressive behavioral capacities. Compared to mice and other rodents, rats have been the exception, as many laboratory tasks of vision have been optimized in rats for over a hundred years^51,52^. Our study represents a fundamental step toward bridging the gap between the rich behavioral history of rats and a more mechanistic understanding of their visual physiology.

## Methods

### Behavior

#### Subjects

All experimental procedures were reviewed and approved by the Harvard Institutional Animal Care and Use Committee (IACUC), under protocol 27-22. All experiments were performed at Harvard University. Animals in this study were female Long Evans rats, 3 months or older, weighing 250-350g (Charles River Laboratories). Rats were housed on a ventilated rack under a reverse 12 hour light:dark cycle with food and water *ad libitum*, except when water-restricted for behavior training. A total of N=56 rats were trained on the basic three-port task, out of which 48 passed criterion. A subset of these rats are part of a previously published dataset^17^. 36 rats were separately trained on the basic two-choice task to test the reliability of the behavior box and paradigm. For identity-preserving transformations or identity-changing transformations, a subset of 13 trained rats each were tested on novel images. Rats were kept on a water schedule in which they received the majority of their water during behavior training sessions. Rats were given *ad libitum* water for 1 hour if few (<100) trials were performed.

#### OpenRatBox

All animals were trained using a custom low-cost, high-throughput, modular training system, OpenRatBox. The frame of the box was composed of custom-cut aluminum extrusions (80/20, Inc.). For the present study, four training boxes were vertically stacked per tower, for a total of four towers. A clear, plastic housing cage snapped and locked into the same position in the behavior box across sessions. A small hole (∼30cm diameter) at one end of the cage allowed the animal to access the monitor and response ports. A custom acrylic mount held three feeding tubes, ∼1 cm apart, (Cadence 7909, 14G Straight Feeding Tube w/ 4mm diameter ball) on an aluminum frame (MicroRax) positioned directly in front of the front access hole of the cage. Each port was coupled to a capacitive sensor. The capacitive sensors (Phidget Touch Sensor 1129, Calgary, Alberta, Canada) were controlled by a USB microcontroller (Phidget Interface Kit 1018). The two flanking ports, which served as the reward ports, were connected to a syringe pump system (NE-500, New Era Pump Systems, Inc., Farmingdale, NY). The syringe pumps were connected via an RS232 adapter (Startech RS-232/422/485 Serial over IP Ethernet Device Server, Lockbourne, OH). Settings for the syringe port were tested for precise and consistent reward delivery of ∼0.02-0.06 mL per trial, at 0.02 mL increments.

Each box was illuminated with red LEDs and monitored via a USB webcam. Each behavior box was equipped with a monitor (Dell P190S, Round Rock, TX; Samsung 943-BT, Seoul, South Korea), positioned ∼30cm from the animal’s head, and a computer (MacMini 6, OSX 10.9.5 or MacMini 7, OSX El Capitan 10.11.13, Apple, Cupertino, CA) mounted above the main vestibule. Training box computers ran an MWorks server and I/O applications for reward delivery (Arduino, Phidget), while the main control computer ran the corresponding MWorks client to control each box (MWorks 0.5.dev [d7c9069] or 0.6 [c186e7], The MWorks Project https://mworks.github.io/).

#### Visual stimuli

Stimuli were produced following Zoccolan et al.^11^. Visual objects were renderings of three-dimensional models built using a ray tracer package, POV-Ray (http://www.povray.org). Each object was defined as a particular configuration and blend of three starting spheres. Objects were rendered with the same light source location and matched to have approximately equal height, width, and area, as defined by a bounding box surrounding each object rendering. Object transformations (*e*.*g*., size, in-depth rotation) were generated using custom Python wrappers and the POV-Ray API. Morphs were generated by gradually adjusting the relative proportions of each object: the composite spheres defining one object were parametrically shifted into the spheres defining the other. We used the Euclidean distance in pixel space to quantify the difference between each neighboring pair of images. In total, 2,000 morphs were generated, from which 22 morphs were sub-sampled such that the Euclidean distance between successive morphs were equal.

#### Training procedure

The basic task was designed according to Zoccolan et al.^11^. Animals were trained to initiate a trial by licking a center port (feeding tube wired to a capacitive sensor), which triggered the appearance of one stimulus. Animals indicated which of the two objects was present by licking the left or the right port.

##### Handling and habituation

Long Evans female rats (Charles River Laboratories, Wilmington, MA) of about 250 g were allowed to acclimate to the colony environment for about a week after arriving. Rats were habituated to human interaction for 1-2 days, then introduced to the training cages and acoustic signals generated by the behavior rigs. Water-deprived rats were encouraged to poke their heads out of the access hole by manually offering water with syringes connected to a feeding needle identical to the ones used in the behavior rigs.

##### Phase 0

On the first 1-2 days in the training boxes, a glob of peanut butter or Nutella was placed on each of the 3 ports to entice water-deprived animals to poke their heads out of the access hole and engage with the lick ports. The reward ports were triggered to always dispense a small water reward (0.02 mL) anytime the animal licked the feeding tubes. Correct trials were rewarded with additional water, but negative feedback was withheld. This period typically lasted 1 session or less, as animals readily licked the reward ports dispensing water.

##### Phase 1

Animals were trained on a single, default training view of object A and object B (40° of visual angle, 0° in-depth rotation). Animals triggered a trial by licking the center port, after which a stimulus appeared on the screen. Animals had 350 ms to 3.5 s to indicate whether object A or object B was on the screen by licking one of the two flanking ports. To prevent spurious licking, trials were aborted if the animal licked <350 ms from stimulus onset. Correct responses were rewarded with water, and the stimulus was left on the screen for an additional 4 s while the animal licked the reward. Incorrect responses were punished with a 1-3 s negative feedback sequence and time-out period. Negative feedback consisted of a short, high-frequency tone and a black-to-middle gray flicker at 5Hz.

To address response bias, the training protocol tracked whether animals licked the same reward port too many times incorrectly, independent of the stimulus shown. In addition, to prevent response bias due to stimulus presentation order, a limit was set on the number of times the same stimulus could appear in a row (N=5 back-to-back presentations of the same stimulus). If either bias was flagged, the other stimulus was shown until the animal correctly responded, at which point the bias counts restarted. Rats usually took about 3-10 days to achieve criterion performance (70% accuracy).

##### Phases 2-4

For testing identity-preserving transformations, an intermediate set of phases was introduced in which feedback was provided for a subset of new views. In a given session, either changes in size or changes in depth-rotation were introduced, but not both (Phases 2 and 3). A staircase procedure was used to slowly introduce increasingly different views (difference from 0° rotation or difference from 40° size). Rats took about 2-7 days total (1-3 days for size, 1-4 days for rotation, of which there were more levels than size) for the staircase to reach the most extreme views for each transformation axis, while maintaining criterion levels of 70% overall accuracy. After each transformation axis was tested separately, rats that maintained criterion performance were tested for one session on both size and rotation training views (Phase 4, see gray cross in **Figure 1I**).

#### Testing

In the last phase of the paradigm, animals were tested on either identity-preserving transformations or identity-changing transformations.

##### Identity-Preserving Transformations

Rats were tested on the same objects presented at various combinations of size and in-depth rotation. No feedback was provided for a subset of views (test views) in order to assess the extent to which generalization occurred spontaneously. With feedback, animals could learn each stimulus-response mapping for all tested views, whereas true generalization refers to recognition of the object presented at novel, untrained views (as in real-world scenarios). Different animals were assigned different views for the no-feedback condition. Poor performance on test views could be due to poor generalization, or more difficult views, such as smaller sizes. To control for this, an additional subset of views that were matched in size (acuity-matched views) were assigned to each corresponding set of test views.

##### Identity-Changing Transformations

Rats were tested on objects that changed identity across a parametrically-varying morph axis between the two original objects. Morphs were presented at the default view (size 0° of visual angle, 0° in-depth rotation). No feedback was provided on morph probe trials, which were interleaved among the regular trials (<15% of trials).

Responses to morph trials were fit with a logistic function via maximum likelihood estimation^171^ using a Python implementation of psignifit^88,172,173^. We estimated a point of subjective equality (PSE), defined as the morph level at which rats were equally likely to report object A or B. We also estimated a ‘just-noticeable-difference’ (JND) for each animal, defined as half the difference in morph levels when ‘B’ is selected 25% and 75% of the time.

### Surgery

#### Head plate implantation

Aseptic surgical technique was followed during all survival surgeries. A head plate and cranial window were implanted in the same surgery as viral injections using methods modified from mouse cranial window procedures^174^. Rats were administered dexamethasone (2 mg/kg) ∼3 hours prior to surgery in order to reduce brain swelling. Rats were anesthetized using isofluorane in 100% O2 (induction, 3-5%; maintenance, 1.5-2%), and placed in a stereotaxic apparatus (Knopf Instruments, Angle Two, Leica). Eyes were protected from drying out with an ophthalmic ointment (Puralube), and then covered with surgical drape that had a hole cut to expose only the top of the animal’s head. Heart rate, breathing rate, oxygen saturation, and body temperature were measured with a pulse oximeter and commercially available software (PulseOx, Mouseox). Body temperature was maintained at 38°C with a feedback-controlled heating pad.

The top of the head was shaved above the incision site, followed by 1 application of Nair (Church & Dwight Co.) applied against the grain for better penetration to clear the site of hair prior to incision. The exposed scalp was cleaned with saline, then wiped with three rounds of alternating Povidone-Iodine and alcohol swabs (Medline) wiped in a center-out fashion over the skin. A small lidocaine block (<0.5 cc) was administered along the incision site, which spanned from just behind the ears to the back of the head. Using a sterile scalpel, an incision was made down the top of the animal’s head, starting from just behind the eyes to the ears, along the lidocaine block. Using forceps and the scalpel edge, tissue covering the skull was carefully scraped off and toward the wound margin to clear the skull on top of the head, with Bregma and Lambda marks clearly accessible.

Once the skull was exposed, a sequence of steps was taken to treat the skull surface for strong adhesion that would prevent rats from ripping off implants. First, a series of small indentations were placed using a small drill all across the cleaned skull to increase surface area and texturize the skull in preparation for adhesives. Next, an extremely thorough cleaning was done of the bone surface with 2-3 rounds of hydrogen peroxide (Swan) and saline washes, followed by 1 round of 10% citric acid and 3% ferric chloride (Dentin Activator, Parkell S393) for 30 sec that was then thoroughly rinsed with sterile saline). The skull surface was completely dried off with highly absorbent, sterile eye-spears (Medline). The tissue around the wound margin was sealed with Vetbond (3M) to ensure no moisture would leak into any part of the exposed skull where the glue would be applied.

After cleaning the skull, the center of the craniotomy was marked at -7.0 to -8.5 mm AP, 4.5 to 6.5 mm ML, depending on the areas being targeted for each animal. Then, the first layer of adhesive was a thin layer of dental glue, evenly applied in one layer across the exposed skull (Quick Base, S398, Catalyst, S371, and Powder, S396, part of C&B Metabond kit, Parkell S380, all mixed on top of an ice tray). The head plate was placed at 30-40 degrees relative to Bregma, which matched the orientation of the imaging plane and captured most of the targeted areas of visual cortex.

The custom titanium head plate was attached to the skull over the right hemisphere. An adaptor mounted to the stereotax frame held the head plate in place, while the dental glue was carefully applied, being careful to leave the craniotomy site clear. This initial gluing was followed by a bulk gluing of a thicker dental glue (Dentsply Integrity Caulk) that provided structural filling and additional support for the angled head plate. Finally, all remaining gaps around the implant were filled with C&B Metabond dental glue. The implant procedure did not require any bone screws or additional supplements to keep the implant stable across months.

#### Cranial window

Since the head plate was attached at a steep angle, drilling the craniotomy and the remainder of the window surgery could be done on a flat surface by affixing the animal by the head plate. A 4-5mm diameter craniotomy was performed at the marked site by careful thinning of the bone with a dental drill within the circular area (Aseptico). Care was taken throughout the drilling process to keep the thinned region within the circular boundaries using a pair of surgical calipers (Fine Surgical Tools). Skull thinning was complete once the entire circular region was semi-transparent and blood vessels were clearly visible through the thinned skull.

Once the skull was thinned down, the region was kept immersed in sterile saline for the remainder of the surgery. The remaining thinned bone was removed with laminectomy forceps (Fine Science Tools) by gently detaching the thinned bone from the rest of the skull along the circular edge, then lifting off the disc of thinned bone. The dura was cut open using a beveled 36G needle tip that was bent such that the beveled tip slightly curved away from the needle hole. This was effective for hooking the dura with the curved tip pointing upward to gently lift up the dura enough away from the cortical surface in order to create a small incision point, without risking pressure or punctures to the cortical surface beneath the dura. Flaps of dura were then peeled back with fine forceps or spring scissors to expose the brain surface, and tucked away around the edges of the craniotomy. Intracortical injections were performed after the duratomy while the entire area was submerged in sterile saline (see ‘Viral Injections’).

A window composed of stacked glass coverslips (four to five 4mm, plus one 5mm, Warner Instruments) bound with optical adhesive (Norland No.71) was then placed over the brain surface. Care was taken to ensure that no pieces of the attached dura flaps were underneath the window, but rather held back and away from the exposed brain surface by the stacked glass cylinder. The remaining saline was partially absorbed out with sterile eye-spears, and the craniotomy was sealed with cyanoacrylate glue (Vetbond, 3M) over a thin layer of sterile saline. Post-operative animals were administered buprenorphine (0.01-0.05 mg/kg) and carprofen (5 mg/kg) daily for 5-7 days following the surgery.

#### Viral injections

Intracortical injections were made at multiple sites (∼5-9 sites per cranial window, spaced 0.5-1 mm apart) using a microinjector (NanoFil, World Precision Instruments) fit with a 36G beveled needle (NF36BV-2, WPI). A high-titre solution of viral vector (AAV9-syn-jGCaMP7f-WPRE) was diluted to a final ratio of 2:1 with a 20% mannitol solution (Sigma-Aldrich) to promote diffusion. pGP-AAV-syn-jGCaMP7f-WPRE was a gift from Douglas Kim & GENIE Project (Addgene viral prep #104488-AAV9). Trace amounts of Fast-Green (Sigma-Aldrich) were added for visual confirmation of injected solution in the brain (**Figure S2B)**. A total of ∼500-750nl was injected per site at a constant rate of 10-25nl/min at a depth of 750µm below the surface. The exposed brain surface remained submerged in sterile saline throughout the injections.

### Widefield mapping

#### Animal preparation

Animals with broad fluorescence across the whole window underwent retinotopic mapping. Typically, ∼4 weeks was sufficient time for expression throughout the window. About 20 minutes prior to the mapping session, animals were anesthetized with isoflurane (5% induction, 1-1.5% maintenance) and administered a subcutaneous dose of chlorprothixene (2 mg/kg, Sigma-Aldrich). During the mapping session, animals remained lightly anesthetized with minimal isoflurane (0.5%). Anesthesia levels were tested with the paw-pinch reflex and breathing rate. The left eye facing the monitor was checked between trials to ensure it remained open and clear.

#### Tandem-lens macroscope

We created a tiltable, tandem-lens macroscope^96,97^, composed of a USB 3.0 CCD camera (MantaG033-B, Allied Vision) and 2 Nikon lenses (Nikon, 105-mm and 55-mm). Images were acquired at 25 Hz with 3x3 pixel binning (256x492 pixels, ½’ sensor) using custom Python scripts. Epifluorescence illumination was achieved with a 470 nm LED (Thorlabs) that was filtered and reflected through a filter cube that housed an excitation filter, dichroic mirror, and emission filter (Thorlabs). Green fluorescence or reflected light was collected and passed through the filter cube then focused on the CCD detector.

#### Visual stimuli

Visual stimuli were presented using custom Python scripts on a large LCD monitor (LG, 72’ diagonal). The monitor was centered ∼60 cm in front of the left eye, spanning 109 degrees of visual field along azimuth, 67 degrees along elevation. A periodic stimulus of a bar cycling across the screen^18,97^ was presented to the (left) eye contralateral to the cranial window. The bar subtended 5 degrees of visual angle, and was presented as a white bar sweeping across a black background or an apertured bar containing a sequence of natural scene images, drifting over a gray background. Maps acquired with both bar types were comparable. The bar was presented at 0.13 Hz along the azimuth and elevation axes, for a total of 2 (downward, rightward) or 4 (downward, rightward, leftward, upward) conditions. The selected stimulation frequency was one of a subset tested that avoided frequency ranges of known, non-stimulus-driven, physiological signals (*e*.*g*., heart rate or breathing rate) in the ranges of 0.1 to 0.3 Hz. One trial consisted of 10 cycles of the bar drifting in a given direction, and a total of 4-5 such trials were acquired for each direction. To preserve the speed of the bar between azimuth and elevation conditions, the bar traversed up and down the full extent of the monitor’s width centered along the monitor’s vertical extent for the horizontal condition (bar started and ended off screen).

#### Image processing

Raw fluorescence signals were corrected for slow drift by removing the rolling average of each pixel’s time course. The width of the rolling window was set to 2.5 times the length of the stimulation period to remove slow linear and non-linear trends. For each pixel, the time courses for each trial (10 cycles of the stimulus moving along a given direction) were aligned and averaged for each condition (1 of 4 possible directions). We then performed a Fourier spectral analysis on the averaged time series to get a magnitude and phase value for each pixel at each frequency. The strength of the response to the stimulus was calculated as the ratio of the Fourier magnitude at the frequency of stimulation to the sum of the magnitudes at all other frequencies^18,97,98^.

#### Area segmentation

Retinotopic maps were created by taking the phase values for all pixels in the image and converting them to Cartesian coordinates that matched the linear position of the bar on the monitor to the phase of the stimulus cycle that corresponded to that position. Phase maps were thresholded using an empirically defined magnitude ratio, which was the magnitude ratio value when no bar was present (blank condition), which was high enough to exclude regions of the image outside of the brain. To identify the borders between visual areas, maps of vertical and horizontal retinotopy were combined to calculate a visual field sign map^98,99^. The visual field sign was computed by taking the sine of the difference between the vertical and horizontal retinotopic gradients at each pixel. Sign maps were then filtered and thresholded to reveal key visual areas: areas with mirror representations map to 1 and areas with nonmirror representations map to -1.

Animals with ambiguous retinotopic maps (due to patchy viral expression or unclear reference areas) were excluded from further study. Given the significant size of V1 and consistent targeting of a large portion of V1 from Bregma coordinates (see ‘Surgery’), V1 was the most reliable reference area to use for identifying areas LM and LI based on known electrophysiological maps of rat visual areas^71^. We identified a given visual area by a combination of metrics^98^: it’s relative location to other identified areas, it’s relative size, its visual field sign, and averaged movies of the neural responses, in which the representation of the drifting bar can be seen traveling across the cortical surface (see **Supplemental Video 1 and 2**). For visualization, phase and power maps were smoothed with a Gaussian window (FWHM=50µm) and masked by applying a threshold to the magnitude ratio.

### Habituation to head-fixation

We developed a shaping procedure to habituate rats for multi-hour head-fixed sessions (the longest sessions were ∼5 hours). After recovering from surgery (see ‘Surgery’), rats began habituation sessions. Rats were placed in a transparent red cylinder with an angled cut on one end to make room for the head plate attached to a custom steel arm (see **Figure S2D**). The cylinder was also used as enrichment in their home cages. Over the course of 2-4 sessions, animals were given decreasing doses of sedative (Midazolam, starting at 1mg/kg) and increasingly long habituation sessions (∼30 minutes to ∼3 hours). Prior to each habituation session, rats were briefly anesthetized with isofluorane (induction, 3-5%; maintenance, 1.5-2%), then placed in the head-fixing apparatus. Rats often struggled in several bouts within the first 15-20 minutes of waking up, but then remained quiescent for increasing periods of time for the rest of the session. Although we did not train animals on a task, observable signs indicated that the animals were relatively calm (for example, they groomed periodically throughout the session, accepted water, and ate treats while head-fixed).

### Tilting two-photon microscope

To access posterior-lateral areas of cortex while keeping the animal in a natural position, we tilted the microscope around the animal to the angle matching that of the implanted head plate (**Figure S2C**). The microscope pivoted about the focal point of the objective to virtually any orientation, and the microscope body allowed ample space for the animal platform, while maintaining >180º of unobstructed viewing angle (**Figure S2E-F)**. We designed two light paths for two zoom modes: a standard-scale mode imaged a 500x500µm^2^ area, with an estimated point spread function of ∼1x1x2 µm (FWHM, 920 nm) measured with a 16x/0.8NA Nikon objective, while the large-scale mode imaged a 1x1mm^2^ area also at single-cell, resolution with an estimated point spread function of ∼2x2x12 µm. The large-scale mode captured the majority of a given visual area in the rat brain, and in some cases, multiple areas at the same time. Both the standard- and large-scale modes could be enlarged to 500x1000 µm and 1x2 mm, respectively, by increasing the scan angle in software (ScanImage^175^). In all modes, the microscope supported simultaneous two-channel imaging. We also attached two epifluorescence paths: a green channel that was useful for visualizing dye-filled blood vessels (see ‘Two-photon imaging, Data acquisition’) to find targeted regions in a larger FOV before switching to two-photon mode, and a blue channel that could be used for retinotopic mapping of a portion of the window that was intermediate in size between the full widefield maps of the whole window and the two-photon FOV.

To hold the animal, we used a custom-designed C-shaped bracket (**Figure S2D**). The bracket was formed with a steel arm mounted to a platform on one end, and on the other end, a steel adapter piece cut at an angle matched to the angle of the animal’s head plate, typically 30 degrees from vertical. We designed the steel arm to withstand forces applied by rats during experiments, and to avoid obstructing the animal’s visual field. The arm reached over the animal’s head from behind, and attached to the angled adapter, which in turn, connected to the animal’s head plate. The angled adapter held three stainless steel ball bearings that mated with half-sphere grooves on the animal’s head plate, which allowed precise kinematic re-positioning in order to access the same cells across days. The steel arm and head plate were designed to be modular, allowing the same components to be reproduced across two-photon, widefield, and habituation setups.

Light shielding around the objective was used to block light emitted from the LCD monitor. A stack of O-rings (McMaster-Carr, Buna-N O rings 016-018) glued over the head plate created a dish-like receptacle above the cranial window^174^, and mated with another O-ring attached to a black shroud around the objective. This created both a light- and water-tight seal for the water immersion objective during visual stimulation.

### Two-photon calcium imaging

#### Overview

A battery of stimuli were used to characterize responses in primary and lateral extrastriate areas of rat visual cortex. Briefly, a moving bar stimulus was used to map two-photon retinotopic preferences and register two-photon fields-of-view to widefield maps identified with the same stimulus paradigm. To measure more fine-scaled receptive field properties, a tiling paradigm was used to estimate the position, extent, and shape of receptive fields. To estimate responses to features such as edge orientation or direction of motion, a set of square-wave drifting gratings were used. Finally, to characterize more complex object representation, as tested in the behavioral choices of trained rats, a subset of the same object stimuli were used to measure neuronal responses in naive rats.

#### Visual stimuli

Head-fixed animals passively viewed visual stimuli presented on an LCD monitor (LG 72’ diagonal, 1920x1080 resolution, 60 Hz refresh rate) positioned ∼60 cm in front of the left eye, subtending 109° of visual angle along azimuth and 67° degrees along elevation axes of the visual field contralateral to the cranial window. Stimulus presentation was synced to neural data acquisition for each trial using custom software (MWorks, Python).

##### Drifting bar for coarse retinotopy

A white bar subtending 2° or 5° degrees of visual angle cycled across a black screen at a stimulation frequency of 0.24 Hz or 0.13 Hz (similar maps were acquired with either stimulus combination). Four cardinal directions were tested (downward, upward, leftward, rightward). Each trial consisted of 12 cycles of the stimulus, and 4-6 trials were presented for each of the four conditions, totalling 16-24 trials total. Blank trials were also included to measure baseline fluctuations in spontaneous activity, and provided baseline values for thresholding magnitude levels for phase maps (see ‘Area identification and validation’).

##### Tiled gratings for receptive field mapping

To estimate receptive field size and positions, we adopted a standard mapping protocol^106^. The monitor was subdivided into square tiles, and each position was stimulated for 500ms, followed by a 1s inter-trial interval (ITI). On each trial, a square-wave drifting grating (spatial frequency of 0.25 cycles/deg, and drifting speed of 10 cycles/sec) randomly switched direction every 125ms between the 4 cardinal directions. Since the receptive field mapping used a shorter ITI, we restricted the stimulated position of a given trial to be a minimum of 30 degrees of visual angle away from the position of the previous trial. The size of the square tile was either 5° of visual angle or 10° of visual angle. For the small tile (5 degrees), this resulted in 21 positions along azimuth and 11 positions along elevation, totalling 231 positions. Each position was stimulated a minimum of 10 times total across 5 blocks, with 2 repetitions of each of 231 positions per block. For the larger tile (10 degrees), this resulted in 11 azimuth positions and 6 elevation positions, totalling 66 positions. Each position was stimulated a minimum of 10-20 times total across 2-4 blocks, with 5 repetitions of each of 66 positions per block. In general, the larger tile was better for getting more responses from LI FOVs. For direct comparisons of receptive field properties between areas, only responses measured with the smaller tile was used. In a subset of sessions, both the small and large tiles were tested in the same cells, and aggregate metrics were qualitatively similar (data not shown).

##### Drifting gratings

To measure visual feature tuning, we presented square-wave drifting gratings at 8 directions (0° to 315°, steps of 45°). Gratings were presented at either full screen or within a circular aperture whose size was determined by the average receptive field size of the population recorded in previous localizer sessions. All gratings were also presented at two spatial frequencies (0.5 and 0.1 cycles/deg) to target the low and high end of known visual acuity levels^49^, and at two speeds (10 deg/s and 20 deg/s). This set of stimulus configurations resulted in 64 unique grating stimuli, which were repeated ∼20 times in pseudo-random order across 4 blocks, such that each block contained 5 repetitions of each of the 64 conditions. Gratings were presented on a gray background (luminance-matched) for 500ms, followed by 2s inter-trial intervals of blank gray screens.

##### Objects

A subset of the object stimuli tested on the trained rats were used for two-photon imaging experiments. Each morph (7 intermediate morphs, plus the 2 target objects) was presented at 5 different sizes (10-50° of visual angle, in 10° steps). For each stimulus size, mean luminance was measured with a photometer placed at approximately the same position as the rat’s eye to determine the gray-scale values needed to create full-screen stimuli matched in luminance for each stimulus size and morph. Luminance differences between morphs at a given size were negligible, so one luminance control was assigned for each of the 5 object sizes tested. This resulted in 50 unique conditions, each presented a minimum of 30 times across 6 blocks, with 5 repetitions of each of the 50 conditions per block. Stimuli were presented for 1s, followed by a 2s inter-trial interval.

### Data acquisition

Neural imaging data was collected using a custom-built galvo-resonant scanning two-photon microscope (20 kHz; Cambridge Technologies) and a 0.8 WD/16x water-immersion objective (Nikon, CF 175). A mode-locked Ti:Sapphire laser (80 Mhz, MaiTai-eHP DeepSee, pre-chirped, Spectra-Physics) provided 920 nm excitation for both channels. Emission was collected using green (535/50 nm) and red (610/75 nm) filters (Chroma) simultaneously on two photomultiplier tubes (Hamamatsu, H10770PA-40).

Laser power was controlled by an electro-optic modulator (Pockels cell, Model 350-80-LA, ConOptics, Inc.). Power measured at the object ranged between 30-80 mW, likely an overestimate of actual power reaching the tissue through the cranial window. The beam was first expanded 3x with a telescope configuration of 50 mm and 150 mm lenses (Thorlabs). The expanded beam could then proceed through one of two paths. The first led to a standard, high-resolution imaging mode (minimum 500x500 µm) in which the spot size was expanded an additional 4x to fill the back aperture of the objective. The second path allowed for a larger FOV (minimum 1x1mm), with a 2x beam expansion that slightly under-filled the back aperture.

For functional datasets, single plane images were collected at a rate of 44.65 Hz (512x512 pixels; 1mm x 1mm FOV) using ScanImage^175^ (ScanImage 2016, Vidrio Technologies). For anatomical volumes, a 200 µm z-stack was taken in steps of 20 µm simultaneously for both channels. A total of 100 volumes were taken and averaged for anatomical images. Depth was controlled with a Piezo controller/amplifier (Physik Instrumente, PI E-665-CR). For localizer and functional runs (see below), the FOV was placed 150-300 µm below the bottom layer of cranial window or the surface blood vessels.

Two-photon FOVs were registered to wide-field vasculature maps offline by aligning blood vessels present in both images (see ‘Area identification and validation’). Matching points between the two views were manually selected based on uniquely identifiable junctures, and a transformation matrix warped one image onto the other. A set of target candidate FOVs were identified in each animal with a viable window (clear cellular resolution, identified visual areas).

All sessions included a localizer run. Localizer runs consisted of the moving bar and tiled gratings stimuli, and identified the receptive fields of cells within the selected FOV. For a given FOV, the stimulus centroid for subsequent functional sessions was determined by taking the center of mass across all fit receptive fields (see **Figure S6)**. Stimuli that were not full-field were centered at this position. Only FOVs for which the bounding box around the largest sized stimuli would be fully within the monitor’s bounds were accepted for functional runs. Functional runs consisted of the full range of stimulus types (see ‘Visual Stimuli’).

High-resolution images of the animal’s behavior state were acquired in sync with neural data acquisition using custom Python software. A CCD camera (Manta G-033, Allied Vision, SonyICX414 sensor) with a zoom lens was centered on the animal’s face on the side facing the monitor, illuminated with an IR LED. Images (492x656, 9.9 x 9.9 µm pixels) were acquired at 20Hz. Image frames were converted to mp4 videos to be analyzed using DeepLabCut v2.0^176–178^. A total of 16 different videos (20 frames per video, automated K-means clustering), sampled from 12 different imaging sessions and each of the stimulus experiment types, were used to train the pre-trained network (ResNet50, training size 0.95, batch size 8).

### Image processing

Motion correction, ROI selection, neuropil correction, and trace extraction were done with a custom pipeline written in Matlab and Python. Motion correction used rigid transformations within each FOV using custom Matlab code (Chris Harvey lab, Harvard Medical School).

#### Cell mask identification

For ROI selection, an activity map was created by taking the standard deviation across motion-corrected frames within a movie file for each block of trials, and then taking the maximum projection across all blocks. ROIs were selected manually with a circular mask using a custom Matlab GUI. To remove background calcium signals, we estimated neuropil masks as circular annuli of 11 µm width, with the inner edge at 9 µm beyond the outermost edge of a corresponding cell body and the outer edge extending to 20 µm^179^. Pixels from adjacent cell body masks were excluded from the neuropil masks.

#### Time course extraction and correction

To get raw fluorescent traces for a given mask, the fluorescence intensity of a cell at each time point was computed as the average fluorescence across pixels within the mask. To correct for slow drift effects due to long imaging sessions, a correction procedure was applied. First, a baseline F<sub>0</sub> signal was extracted with a sliding filter (20% percentile of a 30 sec sliding window) for each cell in each movie. For each trace, this drift was subtracted from the raw trace before adding back the mean of the baseline signal as an offset.

To account for neuropil signals that could contaminate the soma trace, neuropil correction was applied as follows 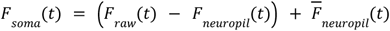 where *t* is time, *r* is an empirical constant defined as the ratio *F*_*bloodvessel*_ /*F*_*neuropil*_(set to 0.7), and 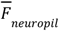 is thetemporally-averaged mean of the neuropil fluorescence^105,179^. Fractional change in fluorescence, ΔF/F(t), following visual stimulus presentation was calculated as: ΔF/F(t)=(F(t)-F<sub>0</sub>)/F<sub>0</sub>, where F(t) is the corrected fluorescence trace during the stimulus presentation, and F<sub>0</sub> is the 1s baseline period prior to stimulus onset. Single response values for a given trial were obtained by averaging the ΔF/F(t) response during the stimulus presentation window.

#### Area identification and validation

Each two-photon FOV was coregistered to widefield retinotopic maps using blood vessel markers. All two-photon imaging sessions began with the acquisition of an anatomical volume, which was a 500µm z-stack taken from the surface. Prior to the start of the imaging session, rats were given subcutaneous injections of Sulforhodamine 101 (SR101, Sigma-Aldrich, S7635) for fluorescent labeling of the blood vessels visible in the red channel. Two-photon blood vessel images were matched to widefield maps offline. Matching points between the two images were selected based on uniquely identifiable blood vessels, then used to identify a transformation matrix to warp one image into the other.

Assignments of two-photon FOVs to visual areas were validated based on retinotopic maps measured with the same cycling bar paradigm used for widefield maps of azimuth and elevation. We applied the same analysis to two-photon FOVs as we did for widefield imaging (see ‘Widefield mapping, Area segmentation’). Phase maps were thresholded by magnitude ratio levels greater than measured in the blank condition (see ‘Two-photon calcium imaging, Visual stimuli, Drifting bar’). Sign maps were obtained from the retinotopic maps with a series of morphological filters^18,20,99^, which were then used to identify patches representing putative visual areas. Two-photon FOVs were segmented based on the visual field sign maps. Since a given FOV could contain more than one visual area, cells were assigned based on both the segmented two-photon sign maps and the wide-field sign maps. Only two-photon FOVs that matched corresponding wide-field maps and had unambiguous sign reversals were included for subsequent analyses.

### Analysis of single cell response properties

#### Estimation of cells with significant visual responses

For each cell, stimulus evoked responses were determined to be significant using a receiver operating characteristic (ROC) analysis from signal detection theory^180–182^. For each cell, a distribution of stimulus responses and a distribution of baseline responses were obtained for each stimulus condition. For each condition, an ROC curve was obtained across a range of criterion levels (50 levels, linearly sampled between the cell’s minimum and maximum stimulus response values) by calculating the proportion of times the stimulus and baseline responses exceeded a given criterion level. The area under the ROC curve (AUC) was calculated for each condition, such that a value of 0.5 corresponds to fully overlapping stimulus and baseline response distributions, while increasingly larger values indicate better separability between the stimulus and baseline distributions. The maximum AUC value for a given cell thus corresponded to the stimulus condition for which the cell’s responses were maximally distinguishable from baseline responses. For each cell, the significance of its maximum AUC value was evaluated by a shuffle test of the baseline and stimulus labels (1000 iterations). Only cells with significant AUCs (*p*<0.05) were included as visually responsive.

For the cycling bar stimulus, the magnitude ratio for each of the 4 conditions was calculated as the ratio of the magnitude at the stimulation frequency to the sum of magnitudes at all other frequencies. A cell was determined to be visually responsive if its average magnitude ratio across elevation and azimuth conditions was greater than an empirically determined threshold. This threshold was estimated by recording neural activity during blank (black screen) trials of the same duration as a stimulation trial, and calculating the magnitude ratio at the stimulation frequency.

#### Estimation of retinotopic preferences in cell bodies and neuropil

##### Retinotopic preferences of background neuropil

The center of each neuropil ring was first assigned a value corresponding to the preferred retinotopic location of the neuropil ring (average over all pixels within the ring). The center was then dilated to a disk of 20µm radius, averaging the preferred retinotopy for overlapping disks. Pixel-wise estimates of retinotopic preference were obtained by smoothing the resulting dilated and averaged image with an isotropic two-dimensional Gaussian filter with standard deviation of 7µm.

For each FOV, the spatial axis corresponding to the direction of maximal retinotopic change for a given retinotopic axis (elevation or azimuth) was identified as follows^105^. First, the two-dimensional pixel-wise gradient was calculated as: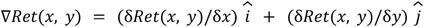

The spatial axis was then computed as the normalized average gradient vector, 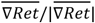, across all pixels. The smoothed neuropil map was then projected onto this mean gradient vector, such that for each pixel, its projected location along this new spatial axis was defined as: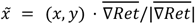. The relationship between a pixel’s preferred retinotopic location (based on the smoothed neuropil maps) and 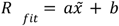 was modeled with a linear function, 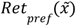, where *a* is the fit parameter (in visual field degrees/cortical µm) that corresponds to the rate of retinotopic progression along the map. The normalized mean gradient and linear fit were computed separately for azimuth and elevation.

##### Estimation of receptive fields

For each ROI, responses at each stimulated location were baseline subtracted (0.5 s before stimulus onset), then averaged across repetitions^106^. An MxN stimulus response map was computed by averaging the response 1s from stimulus onset. The response map *R(a*z,e*l*), where *a*z and e*l* are the retinotopic coordinates in azimuth and elevation, respectively, was then fitted with a two-dimensional Gaussian curve, using the Python implementation of the least-squares Levenberg-Marquardt algorithm^171,183^:

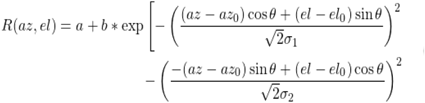

where *(a*z<sub>0</sub>,e*l*<sub>0</sub>)is the center of the 2D Gaussian in azimuth and elevation, σ<sub>1</sub> and σ<sub>2</sub> are the standard deviations along the two axes of the Gaussian, θ is the angle of the Gaussian relative to the azimuth-elevation coordinate system, and *a* and *b* are offset and amplitude parameters, respectively.

The receptive field boundary was considered to be the ellipse defined by the center *(a*z<sub>0</sub>,e*l*<sub>0</sub>) and standard deviations (σ<sub>1</sub>, σ<sub>2</sub>) of the fitted Gaussian:

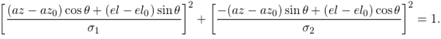

Only fits with R^2^>0.5 and σ between 2.5° and 55° were included for further analyses. To determine whether the fitting procedure yielded a high-quality, reliable fit, we used a bootstrap procedure to estimate confidence intervals (95% CI) for each estimated parameter. Specifically, trials were sampled with replacement, averaged by condition, and fitted according to the procedure described above. This generated a distribution of estimates for each fit parameter, which were then used to determine the 95% CI for each cell’s estimated RF parameters. Only cells with fits lying within the 95% CI (*az*_0_, *el*_0_, *θ*, σ_1_, and σ_2_) were included.

##### Spherical correction

To correct distortions in measured RFs due to presenting stimuli on a flat monitor close to the animal, standard approaches for spherical correcting stimuli were applied in reverse^184^ (see **Figure S4**). First, monitor coordinates were mapped from Cartesian space to spherical coordinates from the rat’s point of view, taking into account the distance, size of the monitor, and angle, relative to the rat’s eye. Typically, the corrective distortion is applied to a visual stimulus to cancel out the distortion caused by the flat monitor covering a large range of visual angle: based on the known measurements between the rat’s eye and the monitor, a 3D model of the monitor can be created, and pixel locations of the monitor are mapped to the spherical coordinates of the monitor. X- and Y-coordinates are treated as angles of azimuth and elevation, and the distortion is applied using interpolation to map horizontal lines to isoelevation lines, and vertical lines to isoazimuth lines. If the stimuli are not corrected to cancel out the distortion, the otherwise distorted percept of the animal can be corrected by applying the corrective distortion directly on the receptive field map. To correct the RF maps, each map was upsampled to match the pixel coordinates of the monitor (resolution was downsampled by a factor of 3 for faster computations). Then, the upsampled map was mapped to spherical coordinates using the 3D model described above (monitor angle=0°, distance=30 cm, monitor width, height, and center, relative to the eye). Finally, the warped image was trimmed back down to stimulus coordinates (degrees of visual angle, in 5° of 10° degree tiles), and the entire RF fitting process was performed on the corrected RF map.

##### Estimation of fine-scale retinotopic scatter

For each FOV, retinotopic scatter was estimated as the deviation, *D*_*VF*_, in degrees of visual field space, from the predicted receptive field center based on its estimated cortical position:

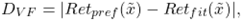

where 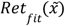 denotes the cell’s measured receptive field center, and 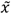 denotes its predicted retinotopic preference according to its projected location 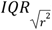 along the mean gradient axis. Cortical scatter, *D*_CX_, in microns, was calculated as the absolute deviation from the predicted cortical position based on the spatial progression defined by the gradient axis:

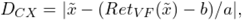

which corresponds to the distance (along the spatial gradient axis) a given cell would have to move along azimuth or elevation in order to form a smoothly progressing retinotopic map.

#### Direction tuning curve fitting

We tested 8 different directions of motion, and two levels of spatial frequency, speed, and size at each direction (a total of 64 distinct drifting gratings). We estimated direction tuning curves using a bootstrap approach combined with conservative criteria for evaluating the stability and goodness-of-fit^105^.

For each cell that exhibited a significant response at a given combination of spatial frequency, size, and speed, a bootstrap analysis (1000 iterations of 20 samples each, to match the measured sample of gratings trials) was performed to compute a mean direction tuning curve. Direction tuning curves were originally sampled in steps of 45°. To obtain a more precise estimate of tuning parameters (preferred orientation and direction), tuning curves were fit with a two-peaked Gaussian^105,185^:

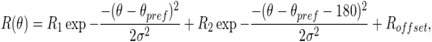

where *R*_θ_is the Δ*F*/*F*(*t*) response for grating direction θ, θ_*pref*_ is the direction evoking the strongest Δ*F*/*F*(*t*) response *R*_1_, *R*_2_ is the amplitude of the second peak at θ_*pref*_+ 180°, and *R*_offset_ is a constant amplitude offset. The model assumes the two peaks of the Gaussian are 180° apart, and that the Gaussians have a common standard deviation, *σ*. Initial parameters were set as follows: *σ* was initialized as the step size (45°), *R*_*offset*_ was the mean of responses to the null directions (all directions except for θ_*pref*_ and θ_*pref*_+ 180°).

To improve the reliability of the fitting and the accuracy of estimated preferred direction and orientation, several additional steps were implemented by following a simplification of the procedure outlined in Liang et al.^105^. First, we added a ninth point at 360° by copying the point at 0° to wrap the input values. Then, the number of input points was increased from 9 to 25 by linearly interpolating the 9-point tuning curve, to more finely sample the curve for the fitting procedure. A bootstrap procedure was used to fit the tuning curves. On each iteration, 20 trials were randomly sampled (with replacement) for each of the 64 conditions, then averaged, interpolated, and fit according to the steps outlined above. The final tuning curves for each cell (one for each unique combination of spatial frequency, size, and speed to which the cell was significantly responsive) were computed from the mean of the fitted parameters across the sampling iterations.

To evaluate the quality of fits, a combination of criteria were used^105^. For each iteration of the fitting procedure, a coefficient of determination, *r*^*2*^, was calculated as the explained variance using least-squares regression to fit the data^171,183^. A combined coefficient of determination, 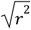, was also calculated for the original tuning curve versus a fitted curve derived from the average of each fit parameter (across the 1000 iterations). These metrics were combined into goodness-of-fit heuristic, *G*_*fit*_:

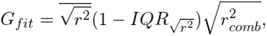

where IQR 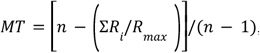 is the interquartile range (difference between the *25*^*th*^- and *75*^*th*^-percentiles of 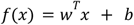 values across iterations. A cell was considered to have a well-fit tuning curve at a given combination of spatial frequency, size, and speed if its *G*_*fit*_ was greater than or equal to 0.5.

##### Axis and direction selectivity

For each cell exhibiting significant responses to gratings (see ‘Estimation of cells with significant visual responses’), a vector sum axis selectivity index (ASI) was computed as a metric for selectivity of motion along a given axis^105,179^. Axis selectivity is distinguished from orientation selectivity, as the latter is typically measured with static gratings, which the current study did not test. To calculate the ASI, the responses for each of the directions was projected onto a circle with 2*i* progression and the magnitude of the normalized vector sum was estimated according to:

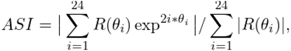

where ASI values ranged from 0 (no selectivity) to 1 (maximum selectivity). Opposite directions were additive, while orthogonal directions canceled each other out. The ASI was computed for 1000 iterations using the same bootstrap procedure as used in calculating the tuning curves. For each cell, the final ASI was computed as the mean ASI across all combinations of spatial frequency, size, and speed for which the cell exhibited significant evoked responses.

A direction selective index (DSI) was computed in a similar way, the responses were projected onto a circle with 1*i* progression:

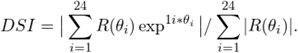

DSI computations were iterated 1000 times, with the final DSI value for a cell taken as the mean DSI across stimulus combinations that evoked significant responses. A cell was determined to be direction-selective if a) it had a significant response to the grating stimuli (see ‘Estimation of cells with significant visual responses’), b) it had a well-fit direction tuning curve (see ‘Direction tuning curve fitting’), and c) it had an average direction selectivity index (DSI) greater than or equal to 0.2^105^ for all stimulus combinations to which it had a significant response. Similarly, a cell was considered to be axis-selective if a) it had a significant response to the grating stimuli, b) it had a well-fit direction tuning curve, and c) its ASI exceeded 0.15 and its DSI was less than 0.2 for all stimulus combinations for which a significant response was observed. For direction-selective cells, preferred direction of motion was determined by taking the circular average of the fitted θ_*pref*_across stimulus combinations (of a given speed, spatial frequency, and size) for which a significant response was observed and direction tuning curves passed goodness-of-fit thresholds.

#### Selectivity and tolerance metrics

Neuronal selectivity to morphs was quantified by a morph tuning index^119,186^, MT=[*n-*(Σ*R*_*i*_/*R*_max_)]/(*n*-1) where *R*_*i*_is a neuron’s response to the *i*th morph, *R*_*max* <*/*_is the maximum response amongst the morphs, and *n* is the number morphs. As a measure of response sparseness, MT ranges from 0 (no shape selectivity) to 1 (maximally shape selective). Size tolerance was quantified by normalizing size tuning curves to their maximum values, then averaging those resulting values that were <1, thatis, *ST*={*R* _*test*_ <*/*_*max*_(*R* _*test*<_)}, where *R* _*test*<_is the mean response to a given test size of a neuron’s most preferred object, and {}denotes the average across tested sizes where *R* _*test*_</_*max*_(*R*_*test*_).

#### Luminance correlations

Since changes in size also change luminance, we estimated the extent to which a cell’s tuning for size could be explained by its tuning for broad luminance. For each cell, its size tuning curve was calculated at its reference morph, defined as the morph eliciting the cell’s maximum response. The cell’s luminance tuning curve was calculated as the cell’s response to each of the size-matched luminance stimuli (fullscreen, grayscale images). The correlation coefficient between the cell’s size tuning curve and luminance tuning curve was taken as a measure for how similar the cell’s responses were for size and size-matched luminance levels. Cells were considered to be luminance-modulated if their size and luminance tuning curves were significantly correlated or anti-correlated (Pearson’s correlation coefficient, *p*<0.05), and were excluded from single neuron characterizations of object selectivity and view tolerance (**Figure S5F**).

#### Single neuron discriminability

Since the animals were naive, a cell’s preference for one object over the other should reflect intrinsic, as opposed to learned, feature selectivities. To quantify discriminability for cells that did exhibit a preference, selectivity for one object over the other was determined with a Mann-Whitney rank test. Only object-selective cells, defined as cells that were significantly selective (*p*<0.05) for one or the other object, were included in the single-neuron discriminability analysis (**Figure S5A-E**).

For these object-selective cells, a receiver operating characteristic (ROC) analysis was used to determine the extent to which the two objects could be discriminated^4,181,182^. Given two response distributions that arise from the two different alternatives (object A or object B), an ROC curve was generated by computing the proportion of trials for alternative 1 (object A) in which the response exceeded criterion versus the proportion of trials for alternative 2 (object B) in which the responses exceeded criterion, for a range of criterion levels. 50 criterion levels were used that linearly spanned the range of the cell’s minimum and maximum responses. The area under the ROC curve (AUC) was taken as a measure of discriminability between the two distributions, that is, how well the neuron could discriminate object A from object B. Cells that passed performance criterion of 70% accuracy on classifying object A and B were subsequently tested on intermediate morphs at their best stimulus size, and responses were fit with a neurometric curve^88^.

### Analysis of population responses

#### Population discriminability

To quantify discriminability, we trained linear classifiers (support vector machines, SVMs) to discriminate the two target objects from the neural activity in each area. The linear-readout scheme represents a biologically plausible processing step that amounts to a thresholded sum taken over weighted synapses^4,122^. Linear support vector machines (SVMs) were trained to discriminate object A from object B from neural responses. Each presentation of an image produces a population response vector **x** of size *N*x1, such that repeated presentations form a cluster of points in *N*-dimensional space (**Figure 5A**, left). The SVM estimates a linear hyperplane that best separates the two classes of responses by maximizing the normalized margin between the two response classes while minimizing classification errors (neural responses are placed on the wrong side of the hyperplane). For a dataset of *N* neurons, the linear readout amounts to:*f*(*x*) = *w*^*T*^ *x*+*b*, where **w** is a *N*x1 vector of the linear weights applied to each of *N* neurons (defines the hyperplane’s orientation), and *b* is a scalar bias term that offsets the hyperplane from the origin. The hyperplane and bias for each classifier was determined by a support vector machine (SVM) using the scikit-learn machine learning library (LinearSVC^187^) with a linear kernel, the SVC algorithm, and cost (C) set to 1.0.

To classify a given image from the population response, a response vector **x** (population response to one image) was applied to the classifier, and negative values of *f*(**x**) indicated object A and positive values indicated object B. The data were split into train and test sets (20%) after balancing numbers of samples per condition. We used a 5-fold cross-validation procedure on the training set to fit and evaluate each model. Test performance was defined as the proportion of correct answers on the held-out test images never included in training.

#### Linear separability and generalization

To test linear separability (**Figure 5C**), 80% of the trials corresponding to object A and object B for each size were combined to train and evaluate the models, while the remaining 20% of trials was used to measure classifier performance. To test generalization, each classifier was trained to classify object A and B at one of the 5 sizes, then tested on each of the remaining 4 untrained sizes. Each training set (for each size) included 80% of the trials for a given size, while the test sets contained either the remaining 20% of trials of the same size (test accuracy on ‘trained’ conditions) or 100% of the trials at one of the other sizes (test accuracy on ‘novel’ conditions).

#### Population sampling

In analyses in which a given metric, *e*.*g*,. classifier accuracy, is presented as a function of the number of neurons in a pseudo-population, we applied a resampling procedure to measure the variability that can be attributed to the particular subpopulation of neurons or subset of trials used for training versus testing. On each iteration, we sampled a new subpopulation of neurons that were randomly selected (without replacement) from all cells aggregated across imaging sites and animals, for a given visual area, and trials were randomly assigned for training and testing (without replacement). Error bars were calculated as the standard deviation (s.d.) of classifier performance across 100 iterations. Chance performance was computed by randomly assigning objects or images associated with each response vector and repeating the classification analysis.

#### Signal and noise correlations

We computed signal and noise correlations in population responses for each imaging site. Signal correlations were computed as the Pearson correlation between the trial-averaged stimulus responses for pairs of neurons. Noise correlations were computed as the Pearson correlation of single-trial responses of a given stimulus condition for a pair of neurons, then averaged over stimuli.

### Quantification and statistical analysis

For all pairwise tests, Wilcoxon signed-rank tests were used, unless otherwise specified. Significance values were set to *p*<0.05(*) or *p*<0.01(**). For comparisons between visual areas, Mann-Whitney U-tests were used. A Benjamini-Hochberg/Yekutieli procedure was used to control the false discovery rate for multiple comparisons^188^.

## Acknowledgements

We would like to thank Zachary Werkhoven, Cindy Poo, Mark Andermann, Cris Niell, Sihao Lu, and Vanessa Ruta for their helpful comments on the manuscript. We are also grateful to Bence Ölveczky, Cindy Poo, Julie Lee, Josh Siegle, and Zachary Werkhoven, for advice and early stage feedback related to this manuscript. We would like to thank Max Joesch for assistance in building the microscope, Ben Vermaercke for helping with initial prototyping at the start of the two-photon imaging efforts, Brett Graham for technical support during the design of the behavior system, and Lily Soucy for the artwork shown in Figure 2. JYR would like to thank Venki Murthy and Mark Andermann for their support from the start of this project to the completion of this manuscript. This work was supported by the Richard A and Susan F Smith Family Foundation and IARPA contract #D16PC00002. CE was supported by the National Science Foundation (NSF) Graduate Research Fellowship Program (GRFP). JYR and JM were supported by the Harvard Brain Science Initiative (HBI), the Center for Brain Science, and the Department of Molecular and Cellular Biology at Harvard.

## Author Contributions

JYR and DDC conceptualized the idea; JYR designed and created the behavior system, as well as the imaging and behavior experiments, with guidance from DDC; JYR analyzed imaging and behavior data; JYR, ES, JG, and DDC developed the two-photon microscope system; JYR and ES developed the widefield tandem-lens macroscope; JYR and DDC wrote software for data acquisition; JYR and CE collected two-photon and widefield imaging data; JYR and CE performed injections and surgeries, and analyzed mapping data; JYR and JM collected behavior data; JYR wrote the manuscript; all authors reviewed the manuscript.

## Supplemental information

See ‘Supplemental Figures and Videos.’

