## Supplemental Figures and Video Legends for "Neural correlates of visual object recognition in rats"

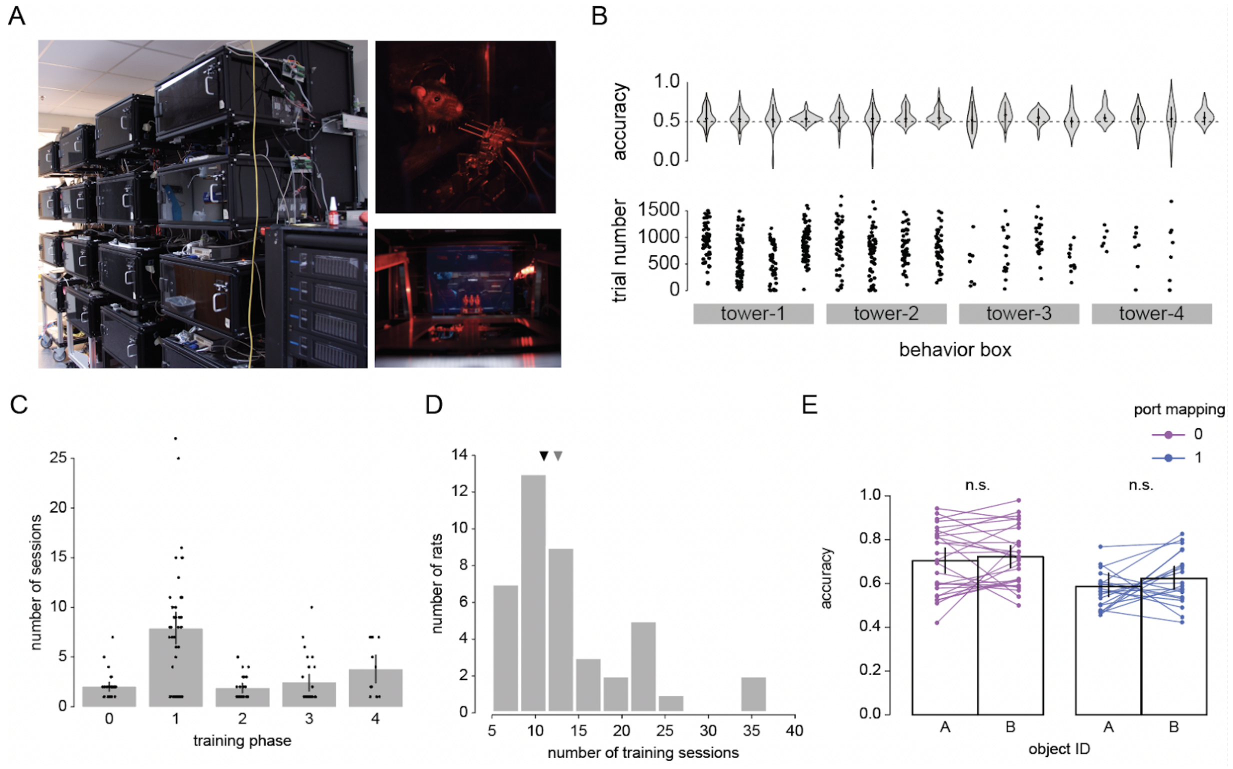


**Figure S1 — Training metrics and validation of high-throughput visual behavior system.** **A**, *Left*: Photograph of four of our towers of stacked OpenRatBox rigs used in the present study. *Right*: Animal looking out of the viewport to access the response ports (top) and the inside view of one box showing metal rails that hold the cage in place and the centered response ports in front of the monitor (bottom). **B**, Top: distribution of accuracy for all animals and sessions, split by behavior box (4 boxes per tower) and tower (4 towers total). Bottom: distribution of number of trials for each animal and session, split by behavior box and tower. **C**. Number of sessions per training phase (N=48/56 rats learned the task). 0: Always Reward, 1: Default View, 2: Size Only, 3: Rotation Only, 4: Size or Rotation Cross (see Methods and Figure 1I). Each dot represents one rat. Bars show mean±SD. **D**, Histogram of the total number of training sessions to reach criterion (across all phases, N=48 rats). Black and gray triangles denote the median and mean, respectively. **E**, Accuracy on the discrimination task, split by object identity (object 1 or 2) and port mapping (whether to lick right for object 1 and lick left for object 2, or vice versa). Each pair of lines represents one rat. Colors indicate arbitrary port assignment for the standard two-choice paradigm (left and right ports mapped to object A and B, respectively, and vice versa).

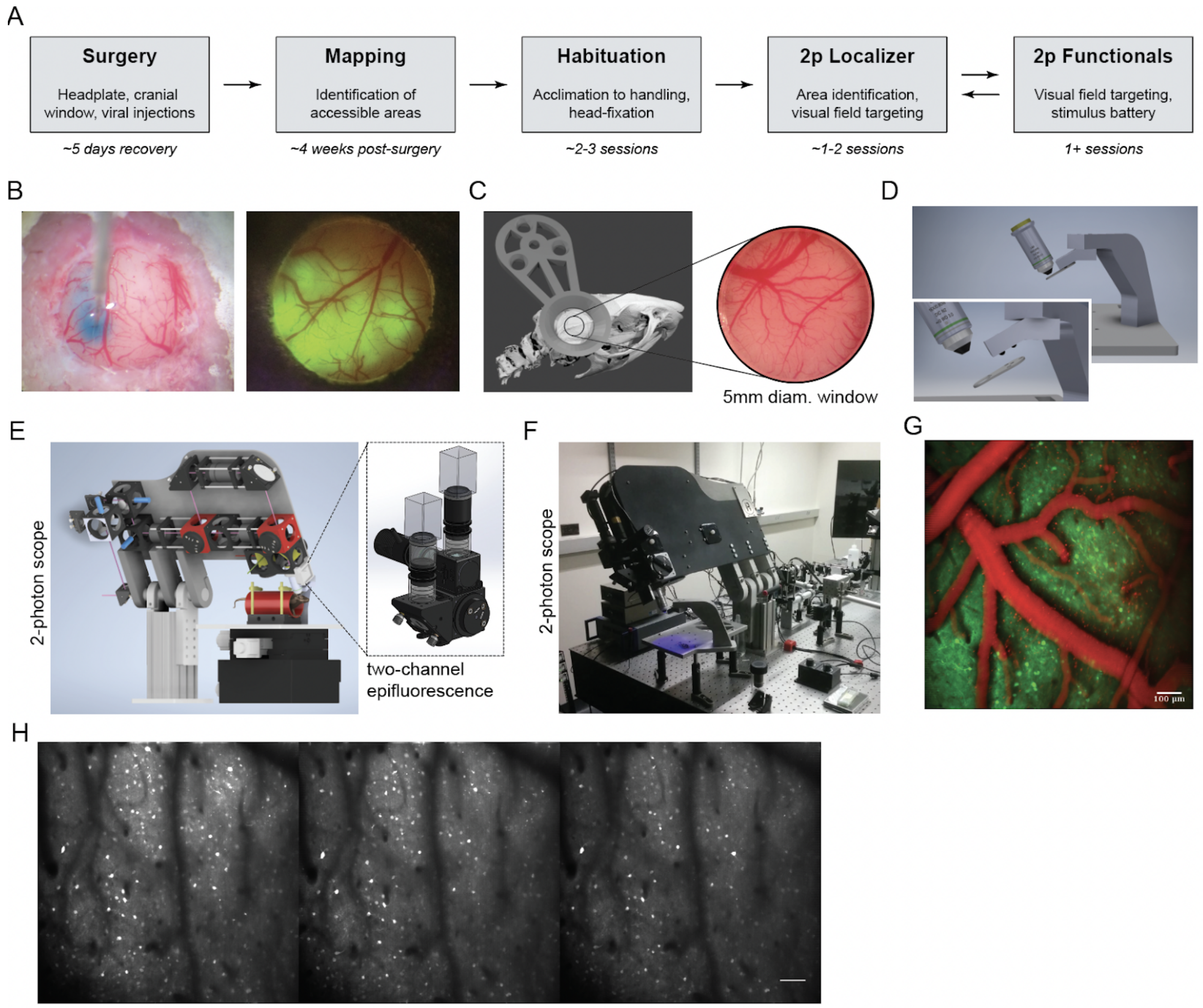


**Figure S2** — **Pipeline for cellular resolution imaging in awake, head-fixed rats.** **A**, Detailed experimental pipeline. 1) Surgery: Animals first undergo chronic implant and cranial window surgery. Viral delivery of the calcium indicator is done during this surgery, as ~4 weeks are needed for expression to come online throughout the window. 2) Mapping: Once most of the window exhibits fluorescence, the mapping procedure identifies which visual areas are identifiable and accessible within the extent of the window. Viable cranial windows must meet several criteria for animals to continue through to the next stage: sufficient expression, identifiable visual areas, and any of the three areas of interest (V1, LM, or LI/LL). 3) Habituation: These animals are then habituated over several days in preparation for two-photon imaging. After habituation, candidate imaging sites, or fields-of-view (FOVs), are selected for two-photon characterization. 4) Localizer: Candidate FOVs are then imaged to validate visual area assignment via retinotopic preference and gradient (see Methods), to target stimulus locations in the visual field based on the receptive field positions of the cells in the FOV. 4) Functional imaging: Vetted FOVs are characterized with a battery of visual stimuli. Localizer runs are also repeated during functional sessions. **B**, *Left*: Exposed cortical surface after craniotomy and durotomy during a microinjection of AAV9-GCaMP7f. Blue, dye used to visualize spread (see Methods). *Right*: Fluorescent view of GCaMP expression in a cranial window. **C**, Schematic of a rat skull with a custom titanium headplate positioned over the approximate location targeted for lateral extrastriate areas in the rat. Circle and inset indicate the position of the optical window implanted over these areas. A photograph of the cortical surface accessible for imaging beneath the 5 mm diameter optical window is shown on the right for illustration. **D**, A thick, custom-cut steel post is affixed to the platform (modular design for both habituation setups and imaging setups). An angled bracket, custom-cut to match the desired angle of the objective, contains 3 grooves that hold spherical ceramic balls (close-up, inset), which mate with the titanium headplate. **E**, Schematic of the microscope housing and optical path along the rotational axis of the microscope. Optical components are included for reference. *Inset*: An epifluorescence microscope attached to the two-photon microscope with dual-channel imaging for FOV identification one magnification step lower than the lowest two-photon zoom. **F**, A photograph of the actual microscope. Beneath the objective is the modular platform and kinematic mounting bracket used to hold the animal. **G**, Dual-channel acquisition for red and green channels. Red, psuedo-colored SR101 blood vessel image. Green, pseudo-colored GCaMP image below the surface. **H**, Example of multi-session imaging of one FOV accessed in three separate days. Each site was identified by eye. Images show the motion-corrected average for each site. No corrections to register one session with the next. Scale bar, 100 µm.

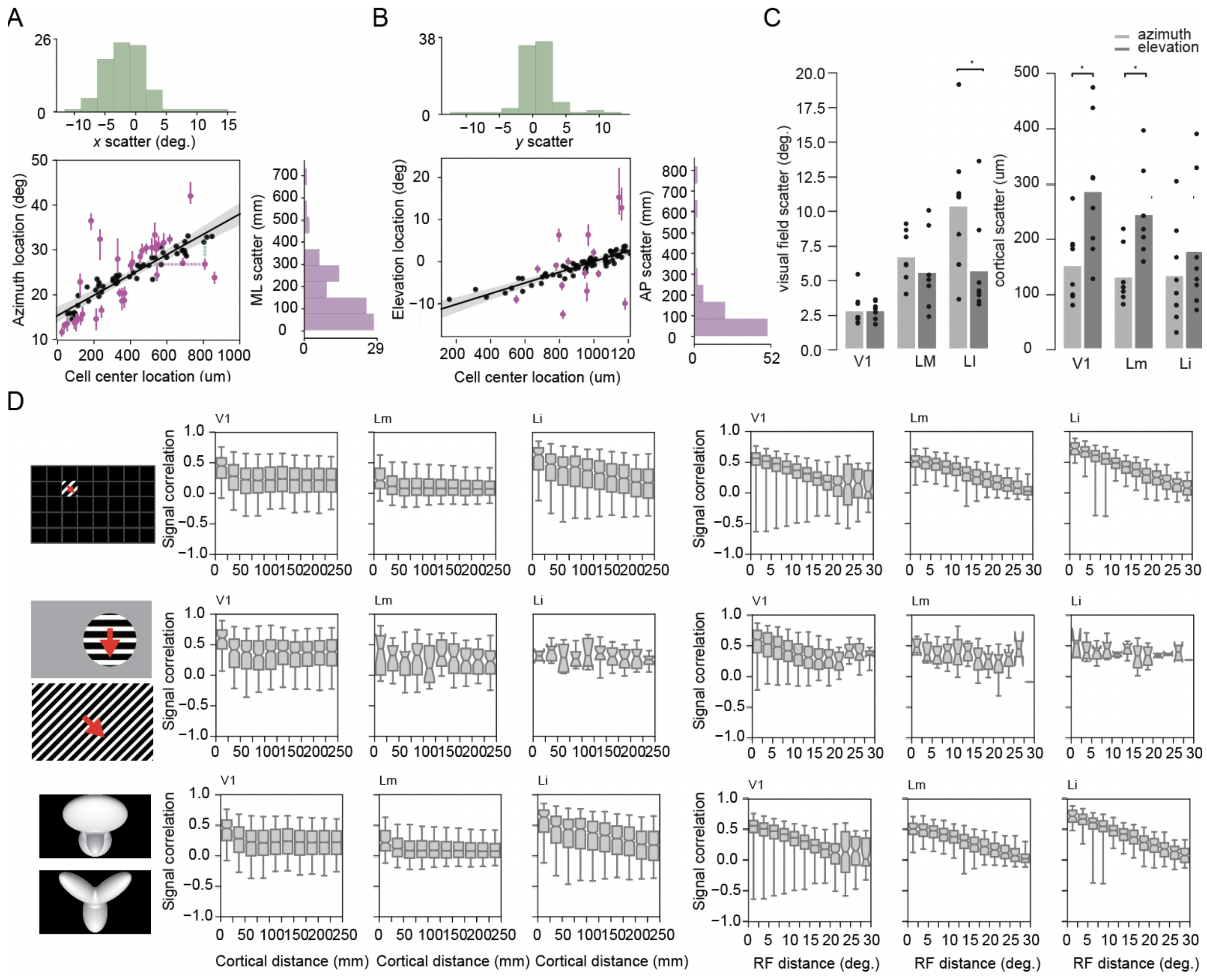


**Figure S3** — **Retinotopy and feature tuning is scattered across cells. A**, Illustration of scatter in a cell's cortical position (purple) and scatter in its receptive field position (green) for an example FOV in V1. Scatter plots show each cell's retinotopic preference versus its cortical location along azimuth. Marginal distributions show histogram of scatter in visual field position (green) and cortical position (purple). Black line, linear fit of the predicted retinotopic gradient (projection of the neuropil map onto the mean gradient vector, see Methods). Shaded bands, 95% CI. Black dots denote cells with reliable receptive field estimates (see Methods). Magenta dots mark "true deviants," or cells with measured retinotopic preferences that significantly deviated from predictions made by a linear model based on cortical location. Magenta lines, 95% CI of each cell's measured and fit retinotopic preference. Green dotted vertical line indicates deviation in preference from the mean retinotopic gradient ("visual field scatter" in degrees of visual angle). Purple dotted horizontal line indicates the distance of a cell from its expected cortical location in the absence of scatter ("cortical distance scatter" in µm). **B**, Same as (A), but for elevation. **C**, Average amount of scatter in visual field position (left) and cortical position (right) for each imaging site recorded in areas V1, LM, and LI. Each dot represents the mean scatter along azimuth (light gray) or elevation (dark gray) for all cells in a given FOV (*: $p<0.05$, Wilcoxon signed-rank test). **D**, Signal correlation as a function of cortical distance (left column) and receptive field center distance (right column) for each stimulus type, depicted in the left icons: tiled gratings used to measure receptive fields (top), drifting grating stimuli (middle), and static object stimuli (bottom). Note that the red arrows for the drifting gratings are for illustrating motion only and were not present in the stimulus. Notches display the 95% CI around the median, upper and lower whiskers indicate the lesser of 75th percentile + 1.5 interquartile range (IQR) or the maximum value and the greater of 25th percentile - 1.5 IQR or the minimum value, respectively.

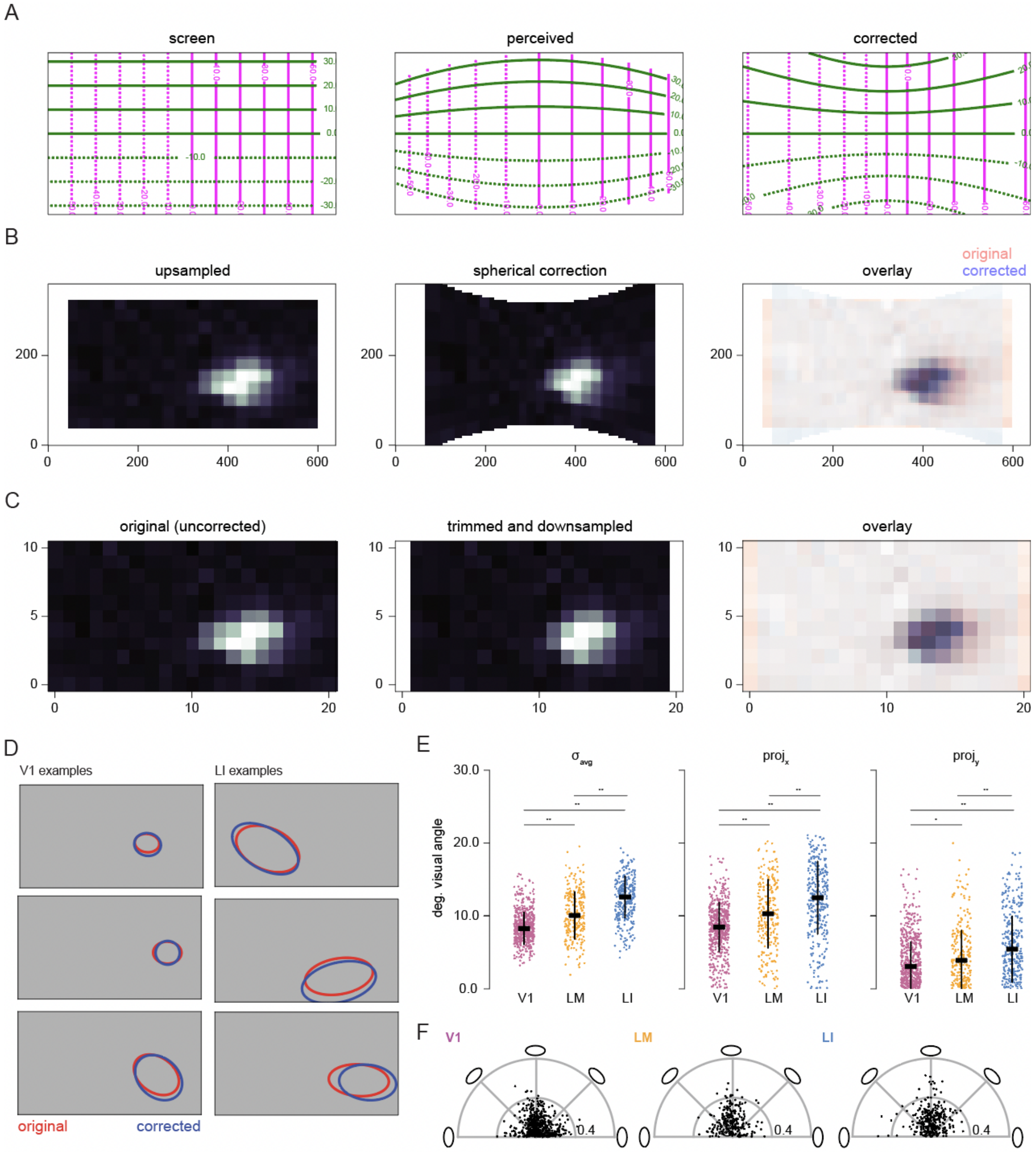
A
screen
perceived
corrected
B
upsampled
spherical correction
overlay
original
corrected
C
original (uncorrected)
trimmed and downsampled
overlay
D
V1 examples
LI examples
original
corrected
E
$\sigma_{avg}$
$proj_x$
$proj_y$
deg. visual angle
V1
LM
LI
F
V1
LM
LI
0.4

**Figure S4 — Spherical correction steps and metrics. A**, *Left*: Screen coordinates of the monitor in degrees of visual angle. *Middle*: Estimate of what the animal perceives without spherical correction[1]. *Right*: Post-hoc transform to warp receptive field responses measured without spherically-corrected stimuli. Green, iso-elevation lines. Magenta, iso-azimuth. Inset numbers denote coordinate space in degrees of visual angle. **B**. Steps for applying a spherical correction to measured responses. *Left*: First, the measured receptive field (RF) response map, corresponding to the cell's average response to each stimulated position, is transformed to pixel coordinates (the response array is upsampled to match the screen size in pixels). *Middle*: Next, the transformation to correct a distorted map is applied to the RF array. *Right*: The overlay shows the warped or corrected RF array (blue) overlaid on the original RF array (red) in pixel coordinates. **C**, Same as (B), but transformed to degrees of visual angle. Each subplot represents the cells' receptive field map corresponding to the correction step depicted above. Units are in monitor locations of the stimulated tile positions (degrees of visual angle), where 0 is the temporal edge (azimuth) or the ventral edge (elevation) of the visual field relative to the animal. **D**, Example receptive field fits before (red) and after (blue) spherical correction for V1 cells (left) and LI cells (right). **E**, Average size (left), measured as half-width at half-max (HWHM) of a double Guassian fit. Also shown is the projection of the major axis of each ellipse onto the horizontal (middle) or vertical (right) axes of visual space for each cell. Each dot represents one cell. Horizontal and vertical bars, mean±SD across cells. **F**, Anisotropy (radius), measured as the ratio of major to minor axes of the fit ellipse, as a function of receptive field angle (theta). Each dot is a cell. Anistropy ranges from 0 to 1, where 0 represents perfectly isotropic receptive fields (a circle), and 1 represents an extremely linear receptive field. Receptive field angles of 0 represent orientations parallel to the horizontal axis, and 90 degrees represents orientation parallel to the vertical axis.

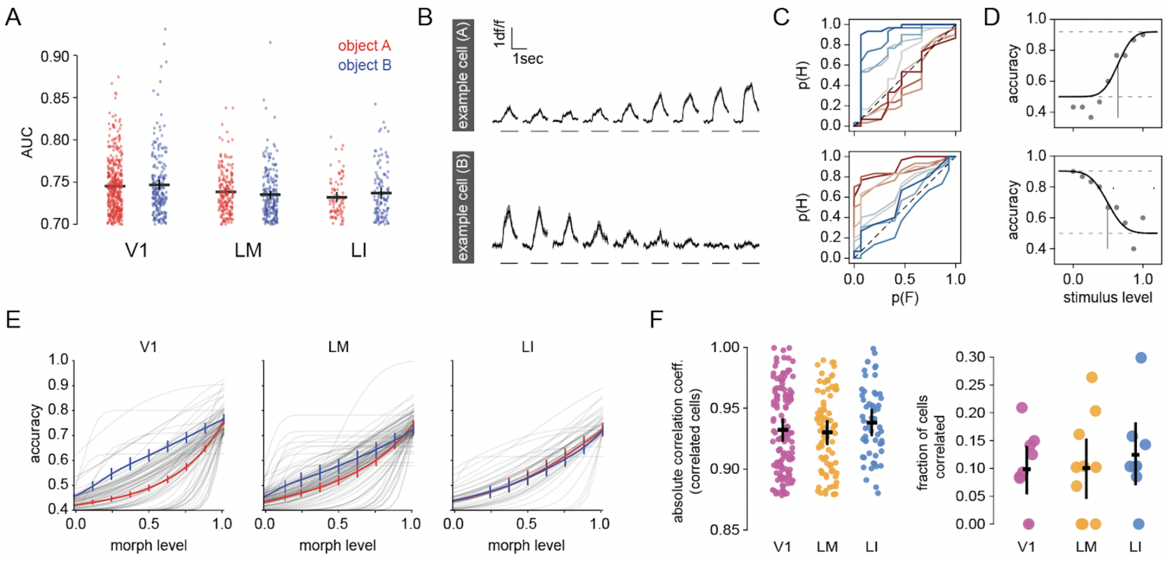


**Figure S5 — Single neuron discriminability. A**, Distribution of discriminability (defined as area-under-the-curve, AUC; see Methods) for cells selective for either object A (red; see example plot in the bottom plot of C) or object B (blue; example plot in the top plot of C). **B**, *Top*: Example time courses at each morph level at the preferred size for a cell preferring object B. *Bottom*: Example time courses for a cell preferring object A. Horizontal bar, stimulus period. Trace and shading, mean±SD across trials. **C**, Top: Example receiver operating characteristic (ROC) curves for the cell shown in the top of (B). The area under these curves (AUC) measures how well a neuron discriminates one image, the preferred object B, from each other image, or the other morph levels. Red corresponds to object A (morph level 0% B) and blue corresponds to object B (morph level 100% B). *Bottom*: ROC curves, but for the cell shown in the bottom of (B). **D**, *Top*: Corresponding neurometric curves fit to the AUC values ("accuracy") at each morph level for the cell shown in the top panels of (B) and (C). *Bottom*: Corresponding neurometric curves for the cell shown in the bottom panels of (B) and (C). **E**, Neurometric curves for all selective, well-fit cells in V1 (left), LM (middle), and LI (right). Only cells that passed criterion performance (70% accuracy) were tested on the intermediate morphs. Response choices were then fit to get a neurometric curve[2]. Thin gray lines correspond to individual cells, only a subset shown for clarity. Red, mean±SD across all individual curves of A-preferring cells. Blue, mean±SD across individual curves of B-preferring cells. **F**, *Left*: Distributions of absolute values of correlation coefficients for all cells with significant relationships between size tuning and luminance tuning (Pearson's, $p<0.05$) for visually responsive cells in V1, LM, and LI. Each dot represents one cell. Vertical and horizontal bars, mean±SD. *Right*: Fraction of cells with significant correlations between size and luminance tuning for each visual area recorded in V1, LM, and LI. Most cells did not have strong size and luminance correlations. Each dot represents one imaging site.

#### Supplemental Videos

**Supplemental video 1.** GCaMP responses to vertical moving bar stimulus for widefield retinotopic mapping.

**Supplemental video 2.** GCaMP responses to horizontal moving bar stimulus for widefield retinotopic mapping.

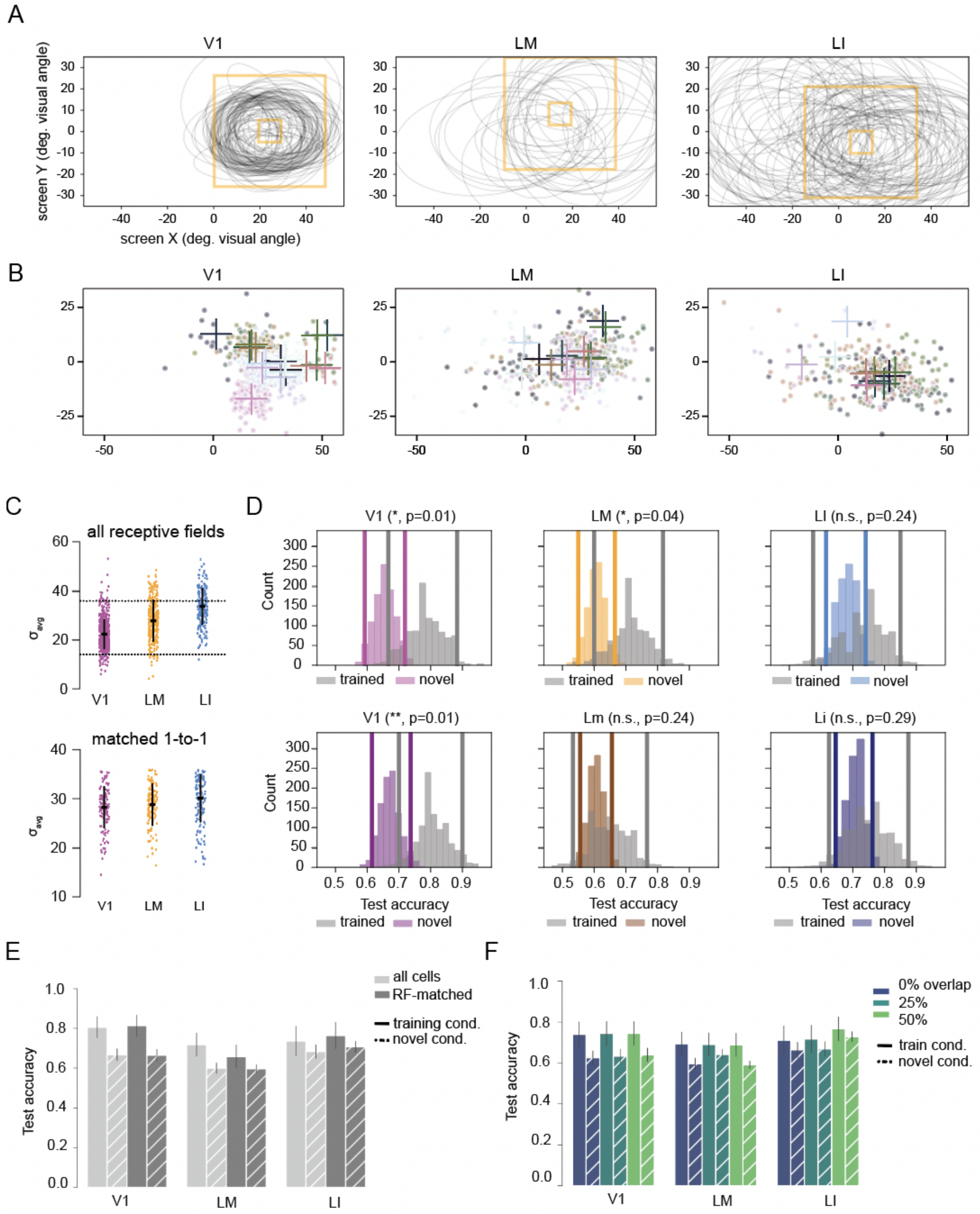


**Figure S6** — **Receptive field size distributions and effect on neural decoding. A**, Receptive fields for an example imaging site in V1 (left), LM (middle), and LI (right). Yellow boxes denote the bounding box around the smallest and largest stimulus sizes for stimuli targeted to the center-of-mass of the receptive fields based on localizer runs (see Methods). Each ellipse represents a fit receptive field for one cell. **B**, Receptive field centers (dots) and corresponding center-of-mass estimates (crosses) plotted in screen coordinates for FOVs recorded in V1 (left), LM (middle), and LI (right). Colors correspond to different FOVs. The FOVs shown here only include cells that were both visually responsive for object stimuli and had well-fit receptive fields in the same session. **C**, *Top*: Distribution of receptive field sizes (average of $\sigma_x$ and $\sigma_y$ from double Gaussian fit, see Method) for V1, LM, and LI. Dotted lines denote the 25th and 75th percentiles of all receptive field sizes. *Bottom*: Distribution of receptive field sizes after sampling within the middle 50th percentile to match all three areas 1-to-1. Crosses represent the median. **D**, *Top:* Bootstrapped distribution of test scores when classifiers were tested on trained (gray) versus the novel (color) stimulus size. Vertical lines represent 95% CI. *p*-values are the fraction of samples that were less than the measured score on the trained stimulus size. *Bottom:* Same as top row, but only including cells from the size-matched receptive field distribution. **E**, Mean±SD of bootstrapped test scores on trained (solid) and novel (hatched) stimulus sizes when all cells were included (light gray) or only cells with1-to-1 size-matched receptive fields across areas (dark gray). **F**, Mean±SD of bootstrapped test scores on trained (solid) and novel (hatched) stimulus sizes when the minimum threshold for areal overlap with any stimulus was 0%, 25% or 50%.

### Supplemental Videos

**Supplemental video 1.** GCaMP responses to vertical moving bar stimulus for widefield retinotopic mapping.

**Supplemental video 2.** GCaMP responses to horizontal downward moving bar stimulus for widefield retinotopic mapping.
